# Ecological constraints foster both extreme viral-host lineage stability and mobile element diversity in a marine community

**DOI:** 10.1101/2025.10.10.681744

**Authors:** Jeffrey Liang, Karine Cahier, Damien Piel, Dario Cueva Granda, David Goudenège, Yannick Labreuche, Laurence Ma, Marc Monot, Charles Bernard, Eduardo P.C. Rocha, Frédérique Le Roux

## Abstract

Phages are typically viewed as very rapidly evolving biological entities. Little is known, however, about whether and how phages can establish long-term genetic stability. We addressed this eco-evolutionary question in an open marine animal associated system, through two longitudinal samplings years apart in an oyster farm, obtaining >1,000 virulent phages and >600 *Vibrio crassostreae* strains. Surprisingly, lineages of phage and bacteria were very persistent, with some phages remaining strictly identical after four years. The phage-vibrio infection network remained modular with multiple lineages co-habitating within individual oysters for long periods of time. Seasonal restriction of *V. crassostreae*, overwintering in wild oysters, and limited viral decay may explain phage stability. Oysters also act as hotspots of activity of diverse mobile genetic element (MGE), hosting plasmids, prophages, phage-plasmids and entirely new classes of satellite-plasmids. Our findings demonstrate how nested ecological constraints can stabilize viral lineages: oysters house vibrio populations that escape host immunity, vibrios restrict phages via receptors and defenses, and satellites parasitize their cognate helper phages. We suggest that ecological context can favor the long-term maintenance of viral lineages and stabilize MGE persistence in the ocean, even amid ongoing mutation and antagonistic co-evolution.

## INTRODUCTION

Viruses are central players in microbial ecosystems, shaping infection dynamics, driving gene flow, and fueling antagonistic coevolution between hosts and parasites^1–4^. Phages are very diverse and their variants have rapid turnovers in communities, fueled by error-prone replication, recombination, frequent horizontal gene transfer (HGT), and antagonistic co-evolution. However, viral populations can be stable over time in natural environments, and the factors involved are poorly understood. Evidence is scarce: for example, a group of *Synechococcus* cyanophages with large genomes showed remarkable genomic stability over at least 15 years in coastal waters (>99% identity in core genes)^5^, even though experimental co-evolution with the host led to rapid diversification in the lab^6^. This contrast suggests that ecological factors—such as host density, range, and environmental conditions—can strongly influence viral evolutionary trajectories. Yet, the simple observation of long-term genomic stability has limited power to reveal the underlying processes, which easier to achieve in model systems where ecological context and evolutionary change can be jointly explored.

Here, we directly address these questions in an open but animal-associated marine environment where bacteria are infected by a large diversity of phages. Oysters are filter feeders, continuously connected to the microbial community in the water column, yet mechanical barriers and innate immunity strongly shape the composition of their resident microbiota^7^. Within this context, we previously identified *Vibrio* populations preferentially associated with oysters^8^. During summer outbreaks, *Vibrio crassostreae* dominates diseased juvenile oysters but is nearly absent from the surrounding seawater. The core-genome phylogeny of *V. crassostreae* revealed eight clades (V1-V8) with varying depths of divergence^9^. Clades V2-V5 and V8 were designated phylopathotypes, as nearly all (>99%) strains in these clades carry the virulence plasmid pGV, in contrast to its much lower prevalence in other clades^10,11^. Phages are also abundant in oyster microbiota^12^ where their predation may influence bacterial dynamics. In a previous time-series study, we isolated *V. crassostreae* and their phages from an oyster farm and performed exhaustive cross-infection assays^9^. We revealed a highly modular infection network, with phage adsorption tightly linked to viral genus-host clade specificity and infection further restricted by intracellular defenses encoded on mobile genetic elements (MGEs). While this provided a detailed snapshot of coevolutionary interactions, it raised key questions: How stable are these bacterial and phage populations over time? And is the modularity of the interaction network conserved?

To answer these questions, we conducted a new time-series sampling four years later at the same oyster farm, generating an unprecedented dataset of over 1,300 phages infecting a single bacterial species, and 604 host genomes. The resulting phage-host infection network again displayed a highly modular structure involving the same *V. crassostreae* clades and phage genera. Remarkably, some virulent phage populations persisted with identical genomes despite the open marine environment. In oysters, temporal patterns of phage genera and host clades reflected dynamic coexistence rather than predator-prey cycles. Beyond these dynamics, our genome analyses revealed an unexpected dimension: oysters are hotspots for HGT and shape the diversity and distribution of MGE in *V. crassostreae.* Plasmids carry genes linked to oyster colonization, while prophages and phage satellites are enriched in defense functions. Our findings reveal a broad diversity of phage-related MGEs with varied architectures and persistence strategies, providing new insights into the eco-evolutionary forces shaping this marine disease system.

## RESULTS

### Stable population of virulent phages within the open marine system

To assess the temporal stability and diversification of lytic phages infecting natural *Vibrio* populations, we resampled the same oyster farm in Brest, France, in 2021, four years after our initial survey in 2017. Across 35 dates between June 28 and September 15, we collected seawater concentrates and oyster plasma and screened them for plaque formation on 153 archival *V. crassostreae* strains representing clades V1-V8. In an initial broad screen, we combined plasma and seawater samples to maximize phage recovery for isolation and sequencing. Phages infecting clade V1 strains were consistently recovered from mid-July through nearly all subsequent sampling dates, whereas those targeting the phylopathotypes V2, V4, V5, and V8 appeared only sporadically (Figure 1). Phages were then selected for sequencing based on host clade and sampling date, without distinguishing their habitat of origin. Over 1,300 isolated lytic phages, 1,033—representing phages targeting all major clades of *V. crassostreae*—were sequenced (Table S1). All sequenced phages belonged to *Caudoviricetes*, with genome sizes ranging from 31.6 to 187.5 kb and encoding 42 to 332 predicted genes.

**Figure 1.**
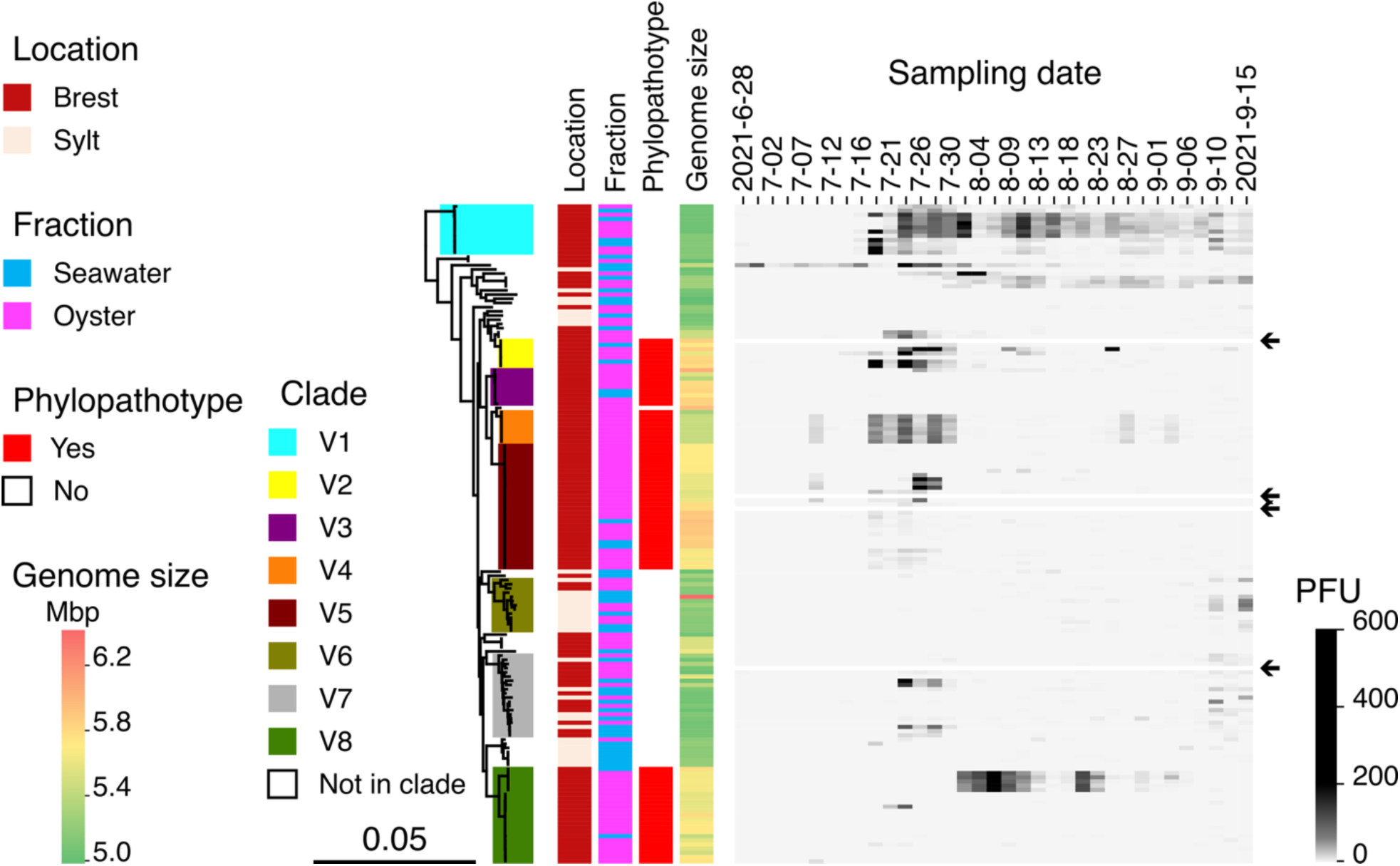
Isolation of lytic phages infecting *Vibrio crassostreae*. During a time-series survey in summer 2021, seawater and oyster plasma were sampled across 35 dates at a single oyster farm in Brest, France. A panel of 153 *V. crassostreae* strains from our archival collection^9^ was used as host “bait” to recover and quantify lytic phages from these environmental samples. Rows represent *V. crassostreae* strains ordered according to a maximum-likelihood core genome phylogeny of 157 isolates, with *V. gigantis* 43_P_281 as the outgroup (based on 2,498 core genes). Arrows indicate strains excluded due to technical issues. Columns indicate the number of plaque-forming units (PFUs) observed per strain and sampling date following infection with 10 μL of seawater viral concentrate (1000x) mixed with 10 μL oyster plasma. PFU counts are shown on a grayscale gradient. One plaque per morphotype and host-date combination was selected for whole-genome sequencing. Additional metadata including clade, strain origin, habitat, genome size are provided.

Using VIRIDIC^13^ we clustered the genomes into 39 genera (>70% identity; LG for Lytic Genera) and 66 species (>95%; LS for Lytic Species) (Figure S1). Classification revealed 899 virulent phages (87% of the total) and 134 predicted temperate phages. Consistent with the predicted temperate lifestyle, members of LG49 and LG50 were also recovered as prophages within *V. crassostreae* genomes (see below). The resulting host-phage isolation network (Figure 2A) showed strong modularity, with clear associations between vibrio clades and specific phage genera—reproducing the cross-infection patterns we found previously^9^.

**Figure 2.**
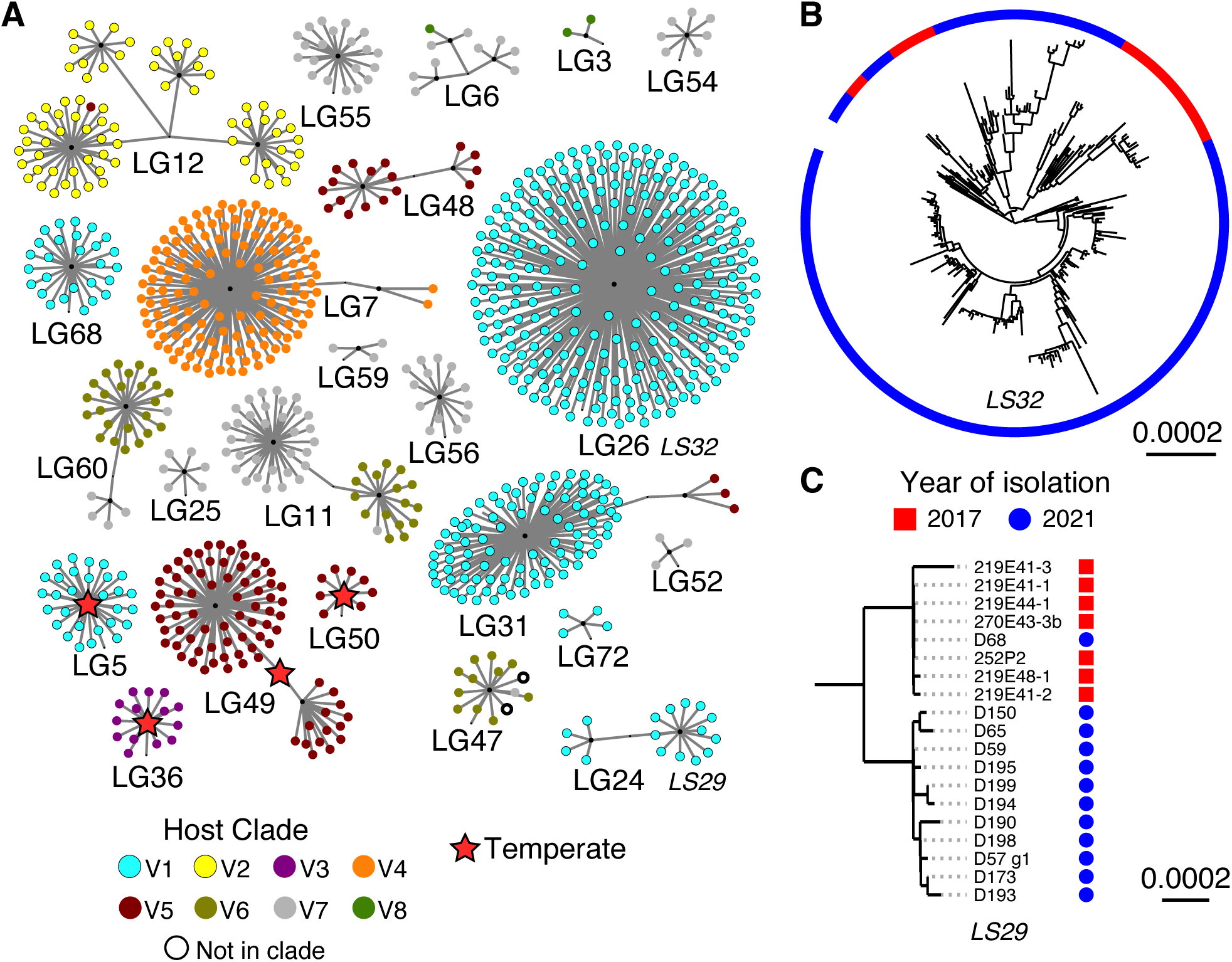
Lytic phages form modular isolation networks and exhibit long-term genetic stability. **A** Hierarchical clustering of lytic phages into genus-level groups (LG) based on intergenomic similarity (VIRIDIC). Each circle represents a phage, colored by the clade of the host strain used for its isolation (same color scheme as in Figure 1). Only genera with ≥3 phages are shown. Most lytic phages (899/1,033) were strictly virulent, but stars mark the four genera predicted as temperate. For LG49 and LG50, this prediction was further confirmed by identifying them as prophages (TG34 and TG89 respectively) within *V. crassostreae* genomes (Figure 5A). Lytic species analyzed in panels B and C (LS32 and LS29) are highlighted in italic. **B-C** Maximum-likelihood whole-genome phylogenies of lytic species LS32 (genus LG26) and LS29 (genus LG24). Scale bars represent substitutions per site (median genome size: 144,041 bp for LS32 and 52,228 bp for LS29). In LS29, one 2021 isolate (D68) was identical to a subset of 2017 isolates, indicating clonal persistence over four years.

Among the 1,033 phages sequenced in 2021, 802 (77%) were from VIRIDIC genera already isolated in 2017. All of these were predicted to be virulent. Thirteen phage species were recovered in both years, with LS29 (7 in 2017, 12 in 2021) and LS32 (28 in 2017, 259 in 2021) especially well represented—enabling direct comparisons across a four-year interval. To assess lineage persistence, we reconstructed time-calibrated phylogenies from whole-genome alignments of LS29 and LS32, correcting for recombination (Figure 2B–C, Figure S2 to S8). In both cases, isolates from 2017 and 2021 were intermingled, rather than forming year-specific clades—indicating long-term persistence of circulating lineages. LS32, a siphovirus (median genome size: 114,026 bp), showed pairwise divergence ranging from 16 to 120 single nucleotide polymorphisms (SNPs) (mean across all pairs: 70.4), with an estimated recombination-free substitution rate of 5.95 × 10⁻⁵ substitutions per site per year. For LS29, a podovirus (52,228 bp), the rate was lower at 1.16 × 10⁻⁵ (95% HPD: 1.55 × 10⁻⁶–2.33 × 10⁻⁵), consistent with marine cyanophages and roughly tenfold lower than dairy siphoviruses^14,15^. These values fall between typical rates of substitution for dsDNA viruses of bacteria and eukaryotes. In LS29, one 2021 isolate (D68) is identical to four isolates from 2017 and differs from the three other 2017 isolates by only 1-6 SNP (Figure 2C), supporting direct lineage continuity and revealing that the same phages can be isolated 4 years apart. Other 2021 isolates diverged by 14-24 SNPs, mostly in a single gene of unknown function (Figure S9), suggestive of rapid host-driven adaptation by homologous recombination in hotspots. These results show that virulent phages persist over multiple years even in an open marine system.

### Stable coexistence of phages and vibrio clades in oysters

We hypothesised that persistence of phage lineages was favored by interactions with co-evolving bacterial hosts. To test this hypothesis, we tracked phage and bacteria populations over time and habitats. To determine the habitat-specific prevalence of phages, we returned to the original samples and separately assayed plaque formation from seawater and plasma against the same hosts (Figure 3A). This analysis revealed contrasting patterns of predation. Phages infecting clade V1 were consistently recovered from both environments, suggesting a persistent environmental reservoir. In contrast, phages targeting oyster-associated phylopathotypes (V2, V3, V5, V8) were almost exclusively found in plasma, with sharp, transient blooms spanning 9-11 days. Although plaque counts were comparable across sources, the 1,000-fold concentration step required for seawater implies that actual phage densities were much higher in plasma, consistent with more intense phage-host interactions within oysters.

**Figure 3.**
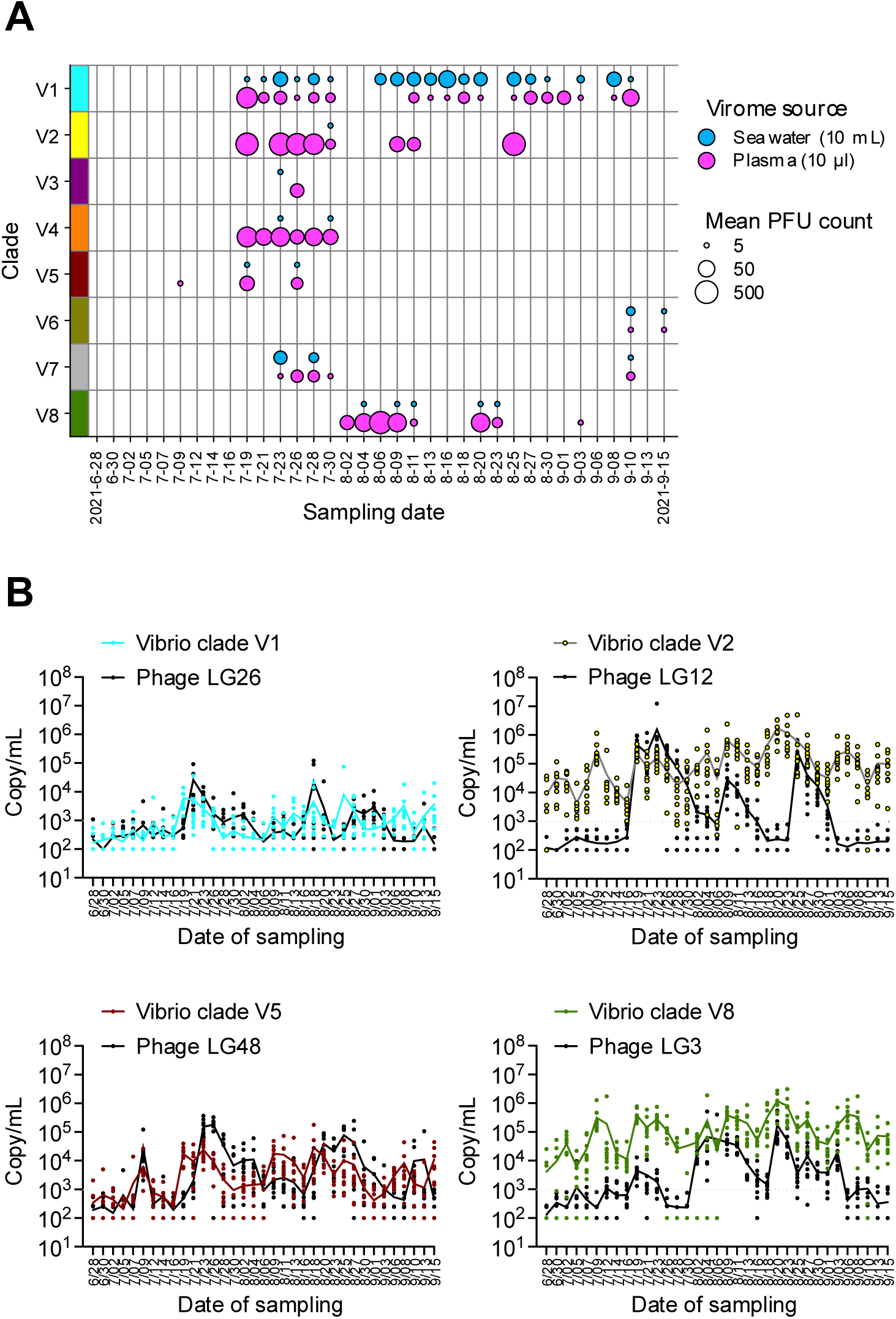
Predator–prey dynamics during the 2021 sampling season. **A** To refine the analysis of temporal and spatial infection dynamics, we separately quantified the average PFU count per clade and sampling date using 10 μL of seawater concentrate (equivalent to 10 mL of raw seawater) or 10 μL of oyster plasma pool. This approach enabled direct comparison of predation pressure across time, habitats and clades. **B** At each sampling date, 10 oysters were collected to quantify specific *V. crassostreae* clades (V1, 2, 5 and 8) and their infecting phages (respectively LG26, 12, 48 and 3) using ddPCR. Each point represents the absolute DNA copy number per mL from a single oyster; lines indicate the geometric mean 202 across the 10 oysters. The dotted horizontal line indicates the limit of detection.

To validate and extend these results using a non-culture-based approach, we quantified the absolute abundance of *V. crassostreae* clades and phage genera in both habitats over time using digital droplet PCR (ddPCR). We first targeted clades V1, V2, V5, and V8 in seawater fractions and in pooled hemolymph from 90 oysters sampled across 35 time points (Figure S10). Clade V1 was rarely detected in either habitat, consistent with its low abundance and the challenge of recovering isolates from this lineage in our collections. In contrast, V2, V8, and to a lesser extent V5 exhibited pronounced fluctuations—spanning up to three orders of magnitude— and reached significantly higher concentrations in oyster hemolymph than in seawater, indicating preferential blooming within the host. We next quantified the abundance of phage genera infecting these clades in the same samples (Figure S11). Phages infecting V2 (LG12), V5 (LG48), and V8 (LG3) showed periodic blooms in plasma, closely tracking host abundance. In contrast, phages infecting the environmental clade V1 (LG24, LG26, LG31) displayed more stable levels across habitats. Together, these data underscore how oyster-associated phylopathotypes and their phages follow a boom-and-bust dynamic within the host, whereas environmental clade V1 and its phages persist through a more stable reservoir.

To examine within-host distributions of both phages and their bacterial hosts across the sampling season, we quantified abundances by ddPCR in ten oysters per time point (Figure 3B). In many cases, peaks in phage and host clade abundance coincided, highlighting the dependence of phages on host presence. However, alpha- and beta-diversity metrics revealed strong variability among individual oysters, with no consistent temporal trends (Figure S12A, B). vibrio clades were positively correlated with one another within individual oysters (Figure S12C), showing that each animal can simultaneously host multiple bacterial clades. On this scale, we found no clear spatial or temporal partitioning of clades or phage genera, preventing conclusions about positive associations or exclusion patterns. Moreover, negative correlations between vibrio clade and phage genus abundance were rare or absent (Figure S12C). Assessing vibrio abundance against contemporaneous, earlier, or later phage samples revealed no detectable predator-prey lags (Figure S13, Table S2) indicating that vibrio dynamics are not tightly coupled to phage replication.

### Heterogeneity in *Vibrio crassostreae* genomes across habitats

The frequent co-occurrence of phages and their bacterial hosts within an individual oyster points to mechanisms—such as resistance or lysogeny—that promote coexistence in oysters. These processes should leave evolutionary signatures in the genome, prompting us to investigate the genetic basis of these dynamics using isolates from the same time series. We analysed 604 sequenced isolates from previous works (157) and 2021 (447, Table S3). The vast majority of the novel isolates originated from oyster hemolymph (95%), reinforcing the idea that the oyster is the preferred habitat of *V. crassostreae*, and are part of the previously described clades V1-V5, V7, and V8 (78%) (Figure 4A, Figure S14). Many of the remaining isolates were from 5 novel clades (V9-V13), among which V9 and V10 are consistently associated with oysters.

**Figure 4.**
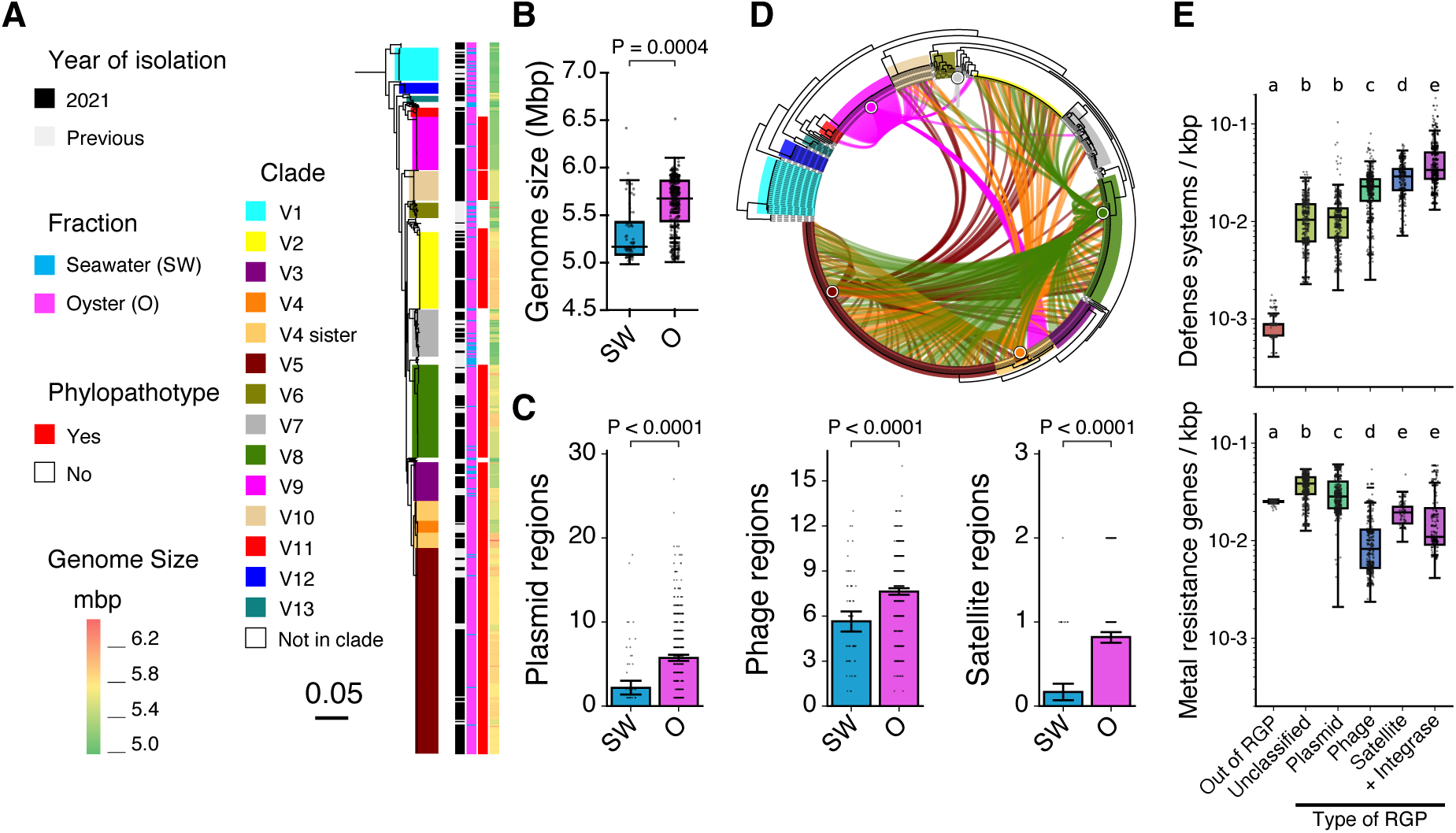
Habitat-specific distribution of MGEs. **A** Rooted maximum-likelihood phylogeny of 604 *V. crassostreae* strains based on 3,099 persistent gene families (outgroup *V. gigantis* pruned). Clades and colors follow ^9^, with new clades (V9-V13, V4-sister) highlighted. Columns indicate year of isolation, sample type, phylopathotype status, and genome size. **B** Oyster-derived isolates have significantly larger genomes than seawater isolates (phylogenetic ANOVA with Holm-Bonferroni post hoc tests, *P* < 0.001). **C** Plasmids, temperate phages, and satellites are more frequent in oyster-associated genomes (phyloGLM with Poisson_GEE regression, *P* < 0.001) **D** Distribution of the virulence plasmid pGV across the phylogeny. Links connect strains to their closest hybrid-assembled plasmid (wGRR > 50%). **E** Mobile genetic elements specialize in different types of cargo genes. Phage defense systems (Padloc) are enriched in genome plasticity regions (RGPs) compared to the core genome, and occur more frequently in phage-, satellite-, and integrase-associated regions than in plasmids. Metal and biocide resistance genes (BacMet2) are significantly more common on plasmids. Significance was tested using Kruskal-Wallis with Dunn’s post-hoc test and Benjamini-Hochberg correction. Colors and letters show compact letter display groups defined by overall similarity.

The genome size in the vibrios range from 5.0 to 6.5 Mbp, with seawater isolates exhibiting significantly smaller genomes (mean: 5.32 Mbp) than those from oysters (mean: 5.62 Mbp) (Figure 4B). The phylogeny-aware analysis of gene exchange rates in the *V. crassostreae* pangenome revealed a species-wide association between core-genome phylogenetic distance and the rates of gene gain/loss (p < 0.0001; Tables S4 and S5). However, this was not the case in V10 (p = 0.964) or in the seawater-associated clades V1 (p = 0.501) and V6 (p = 0.451). In the phylopathotype clade V10, this is likely a reflection of a very recent epidemic expansion while for V1 and V6, this could suggest that clades living in seawater have different pangenome dynamics. Corroborating this, seawater isolates (n = 71) had significantly lower gene turnover than oyster-derived strains (n = 534; p = 0.021), suggesting reduced HGT in the open-water environment.

We identified 32,349 regions of genome plasticity (RGPs) (Table S6), comprising 3,217 plasmid-associated, 4,478 prophage-associated, 450 satellite-associated, and 1,945 integrase-associated regions (following^9^). Plasmids, prophages, and satellites (i.e., parasitic elements that hijack helper phages for propagation) were consistently more abundant in oyster-derived strains (Figure 4C, Table S7). This pattern was particularly striking for the virulence plasmid pGV, which formed a highly reticulated similarity network, indicative of frequent transfer across the species— likely mediated by the oyster-associated phylopathotypes (Figure 4D). We next examined the associations between different MGEs and their genetic cargo. We previously showed that genes encoding anti-phage systems are enriched in RGPs relative to the core genome^9^. Here, we find that these systems are concentrated in phage-, satellite-, and integrase-associated RGPs, but comparatively rare in plasmid-associated regions (Figure 4E, Table S6 and S8). To further characterize plasmid diversity and function, we fully assembled 20 plasmids grouped into five families: the virulence plasmid pGV and four novel plasmids—p1, pMintaka, pAlioth, and pMizar (Tables S9–S13). All encoded conjugation machinery, including Type IV secretion systems and MOB relaxases, consistent with self-transmissibility. Beyond phage defense, plasmids carry functions likely advantageous in the oyster host. For instance, pGV encodes a Type VI secretion system linked to hemocyte toxicity^11^ and genes potentially involved in copper resistance; p1 harbors a metallophore biosynthesis and transport system; and pMintaka encodes microcin synthesis and secretion genes. Compared to other MGEs, plasmids were enriched in functions related to oyster adaptation, including metal and biocide resistance (Figure 4E, Table S6)—traits likely reflecting selective pressures imposed by hemocyte activity, antimicrobial peptides, and the oyster’s tight regulation of metal ions^16^.

### Clade-specificity and episomal persistence of temperate phages

To assess the contribution of temperate phages to genome plasticity in *V. crassostreae*, we screened our bacterial isolates with geNomad^17^, identifying ∼1,600 putative prophages spanning three viral classes: *Caudoviricetes* (tailed), *Tectiliviricetes* (tailless), and *Faserviricetes* (filamentous) (Table S14). Controls using long-read sequenced genomes revealed that predictions of *Caudoviricetes* were more consistent (Figure S15), as seen by^18^. We thus retained 562 *Caudoviricetes* prophages larger than 25kbp, of which 525 were chromosomally integrated and 37 (6.6%) were extrachromosomal (Table S15). VIRIDIC clustering grouped these into 36 genera (TG for temperate genus) and 80 species (TS for temperate species). Among them, 36 species were singletons and 12 occurred in hosts not assigned to a clade. Of the remaining 32 phage species, 30 were clade-specific and 14 were found within the genomes of 2-5 vibrio clades. At the genus level, only 7 of 23 non-singleton genera (30.4%) were clade-specific— substantially fewer than for virulent phage genera (21/30, 70%). Thus, bacterial clade specificity is evident at the phage species but not genus level (Figure 5A). We found 20 prophage species persisting in vibrio genomes sampled between 2001 and 2021, eight of which showed complete genomic invariance across all aligned isolates. We could thus compute the substitution rates for the remaining 12 prophages, like we did for the virulent. We observed substitution rates significantly lower for prophages than for virulent phages (mean ∼10⁻⁶ substitutions per site per year) (Figure 5B). The phylogenies of the best-represented species (Figures S16-S27) reveal striking similarities across isolates. For example, in the species TS30 more than 20 elements are identical (Figure 5C). Together, these results point to tighter co-evolution between vibrio clades and temperate phage species, which show limited diversity and appear to persist for years primarily through vertical transmission within clades. In contrast, virulent phage species are more diverse and infect a broader range of strains within a clade, likely reflecting selection for infection breadth to overcome host defenses.

**Figure 5.**
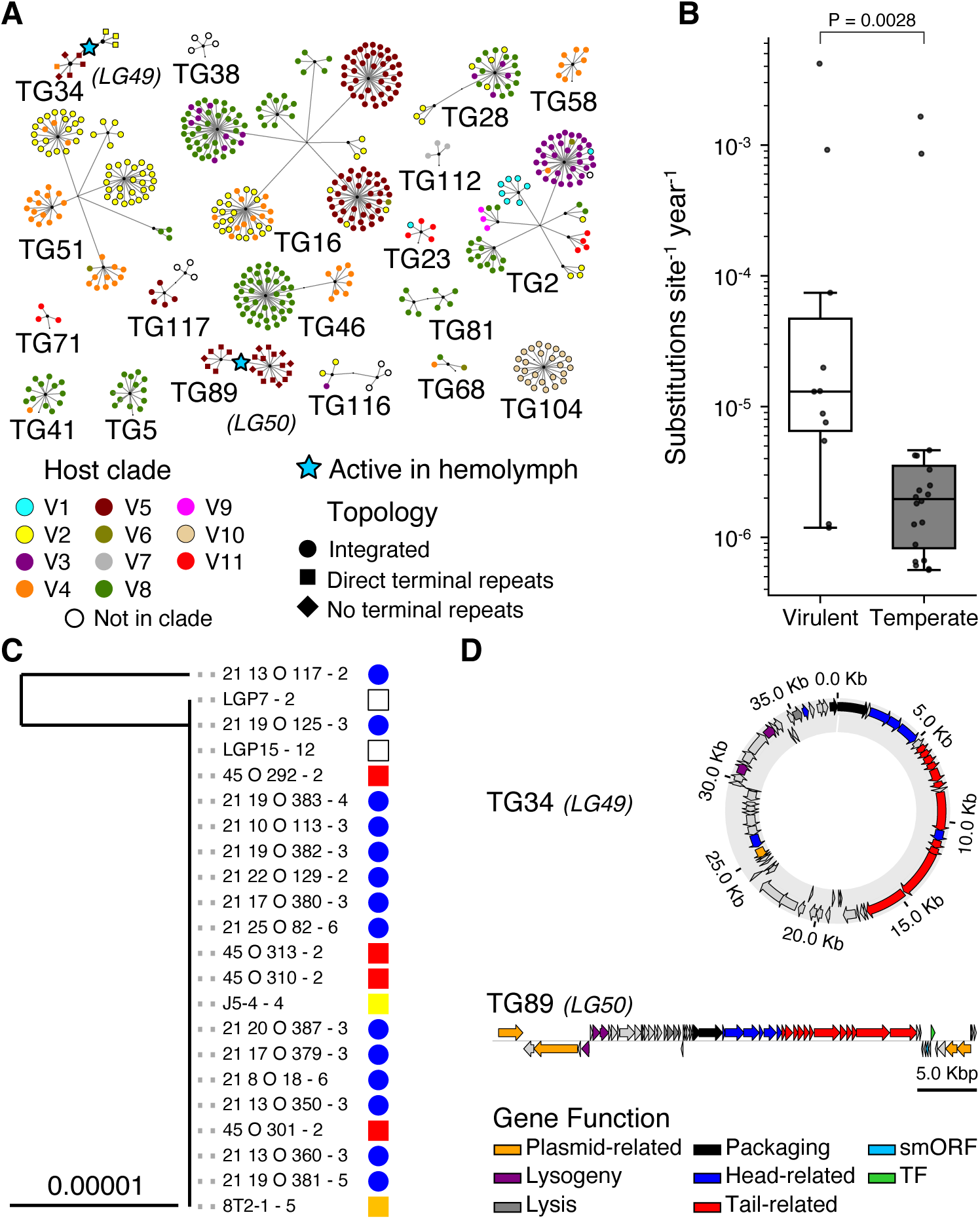
Clade specificity and persistence of temperate phages. **A** Hierarchical clustering of prophages predicted by geNomad across 605 *V. crassostreae* genomes, grouped into temperate genera (TG) using VIRIDIC intergenomic distances. Each node represents a prophage, colored by the clade of its lysogen. Node shapes indicate predicted topology: circles (integrated proviruses), squares (contigs with terminal repeats), and triangles (contigs without terminal repeats). For clarity, only genera with ≥3 representatives are shown. Stars mark TG34 and TG89, two temperate genera corresponding to lytic genera LG49 and LG50 (Figure 2A) that were recovered as active phages in hemolymph. **B** Substitution rates estimated from whole-genome alignments, calibrated with isolation dates (virus for virulent phages; host for prophages), using BEAST v2.6.3 and an exponential relaxed clock. Significance was assessed with a non-parametric Brunner-Munzel test. **C** Maximum-likelihood phylogeny of TS30 shows this prophage persisting in *V. crassostreae* for over 20 years with 100% nucleotide identity. **D** TG34 (circular) and TG89 (linear) are phage-plasmids. Gene annotation revealed both plasmid- and phage-related functions. TG89 also encodes homologs of a transcription factor (TF) and smORF previously described in the linear extrachromosomal phage 63 of *V. cyclitrophicus*^21^.

The two temperate genera (TG34, TG89) correspond to LG49 and LG50, originally identified as lytic phages but predicted temperate by BACPHLIP. Unlike most temperate elements, these stood out because we recovered them as infectious particles in plaque assays, confirming their capacity for lytic activity. They are also unusual in assembling as complete extrachromosomal contigs rather than integrated prophages. Read coverage profiles and ddPCR estimates— showing plasmid-to-chromosome ratios ranging from 3 to 10 depending on the strain—indicate a plasmid-like, low-copy-number replication strategy (Figure S28). Both carry plasmid partition systems (Figure 5D), supporting their identification as active phage-plasmids^19,20^. TG89 is a linear phage-plasmid, maintained extrachromosomally with covalently closed ends, resembling N15 in *E. coli*, VP882 in *V. parahaemolyticus*, and phage 63 in *V. cyclitrophicus*^21,22^. It encodes *telN* (protelomerase) and *repA* (replication initiator), hallmark genes of this phage-plasmid lifestyle. TG89 also carries homologs of a small ORF and transcription factor (smORF and TF in Figure 5D), previously implicated in polylysogeny regulation^22^ though their functional roles in *V. crassostreae* are unknown.

We tracked TG34 and TG89 during the 2021 time series using ddPCR on plasma (free viral particles) and hemolymph (bacteria + viruses). TG89 remained consistently near or below the detection limit across all samples, precluding meaningful ecological interpretation. By contrast, TG34 exhibited sporadic but massive blooms—up to three orders of magnitude increases—in hemolymph, while remaining barely detectable in plasma (Figure S29A). TG34 was the only phage significantly more abundant in hemolymph than plasma, consistent with rapid reinfection and establishment as lysogens rather than long-term persistence in particle form (Figure S29B). These phage-plasmids illustrate an additional dimension of temperate phage ecology: unlike integrated prophages, they persist as an episome and can toggle between lytic activity and lysogeny, highlighting oysters as key environments where temperate phages actively shape both viral and bacterial populations.

Taken together, our results reveal two distinct strategies of temperate phage persistence in the oyster-vibrio system: long-term vertical transmission of clade-specific prophages and episomal maintenance of phage-plasmids capable of switching between lysogeny and lysis. Both strategies underscore the ecological filtering imposed by oysters, which not only stabilizes host-phage associations but also diversifies the ways temperate phages contribute to bacterial genome plasticity and population dynamics.

### Discovery of episomal phage satellites expands the known diversity of MGEs

Two putative prophage genera, TG54 and TG93, assemble as extrachromosomal circular elements with unusually small genome sizes (15.8 and 11.1 kbp) and lack tail genes— suggesting they are phage satellites^23^. Both encode plasmid maintenance functions, including *parA*, *repA*, and a putative *ProQ/FinO*-like regulator, supporting episomal replication (Figure 6).

**Figure 6.**
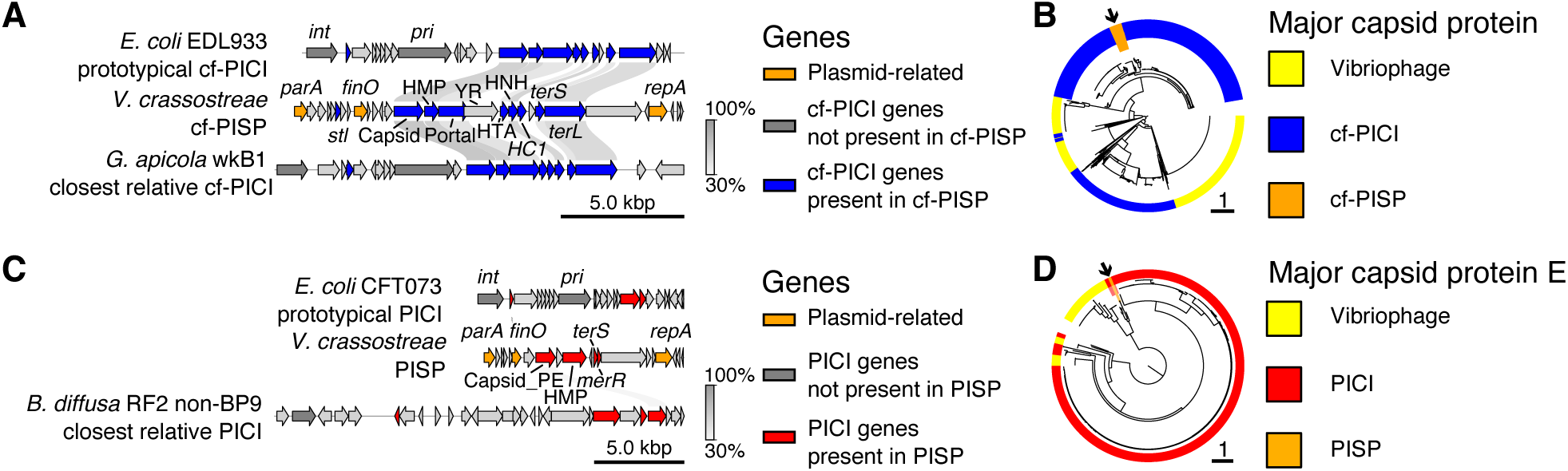
Gene content and phylogenetic placement of novel satellite–plasmid elements. **A** Comparison of gene content between two known capsid-forming PICIs (cf-PICIs) and the episomal element cf-PISP, showing conserved synteny. **B** Maximum-likelihood phylogeny of capsid proteins (Pfam PF05065; n = 1,375) from cf-PISP (n = 30), published cf-PICIs (n = 900), and lytic phages from this study (n = 465). **C** Gene content comparison between two known PICIs and the episomal element PISP, showing divergent gene order but conserved functional modules. **D** Maximum-likelihood phylogeny of PF03864-type capsid proteins (n = 410) from PISP (n = 2), published PICIs (n = 356), and vibriophages from this study (n = 52). In A and C links indicate >30% amino acid identity based on bidirectional best hits (MMseqs). Genes are colored by functional category and presence in satellite-plasmids or predicted substitutes (e.g. alpA, merR, stl). Tree inference was performed with IQtree with Q.pfam+F+I+R substitition model using mafft L-INS-i amino acid sequence alignments.

TG54 was found in 30 vibrio genomes, mostly in the oyster-associated clade V2, isolated both in 2017 and 2021. It was classified by SatelliteFinder as a capsid-forming phage-inducible chromosomal island (cf-PICI)^24^, encoding homologs of structural proteins for head formation (capsid, portal, head maturation protease, head-tail adapter, closure), the HNH endonuclease and terminases for genome packaging, and an stl-like regulator likely controlling induction in response to phage activity. TG54 lacks an integrase but encodes a distinct tyrosine recombinase located between the portal and head-tail adapter genes (Figure 6A). The phylogeny of the capsid protein reveals clustering of TG54 with known Gammaproteobacterial cf-PICIs (Figure 6B) clearly separated from phage proteins, further showing that TG54 is not a satellite and not a cryptic phage. We propose that TG54 represents a satellite-plasmid: an element maintained vertically like a plasmid but transmitted horizontally by hijacking helper phage particles. Because it is not a chromosomal island, we replace ‘island’ in cf-PICI with ‘satellite-plasmid,’ naming this new element a cf-PISP (capsid-forming phage-inducible satellite-plasmid). A search of public databases using its capsid and ParA protein sequences identified similar elements in at least two other *Vibrio* species, but none outside the genus (Figure S30).

TG93, detected in two strains from clades V4 and V8 in 2021, carries fewer genes than cf-PISPs and was not flagged as a satellite by SatelliteFinder. Manual inspection, however, revealed divergent homologs of hallmark genes from phage-inducible chromosomal islands (PICIs), another class of phage satellites^25^ (Figure 6C). The element encodes Capsid protein E and a head maturation protease, together with a small terminase and merR-like transcriptional regulator. We did not detect an integrase in TG94; the gene may be dispensable in this element because it is a plasmid. Based on its gene content, we suggest that this element shares the PICI strategy of head assembly disruption, interfering with capsid subunit assembly to form a physically smaller prohead that excludes the larger helper phage genome and preferentially packages the satellite genome. The capsid protein sequence clusters with satellite-encoded and not phage-encoded capsid proteins in the phylogeny (Figure 6D), supporting our identification of this element as a second type of episomal satellite-plasmid. By analogy with cf-PISP, we name this smaller element PISP (phage-inducible satellite-plasmid). Related elements were detected in three other *Vibrio* species but not in any other bacterial genera (Figure S30).

## DISCUSSION

Microbial populations are often seen as highly dynamic, with rapid turnover and diversification, yet phages—despite being considered the fastest-evolving biological entities of these populations—sometimes persist stably for years in natural environments. The oyster-vibrio-phage-satellite assemblage that we study showed it is a powerful model for disentangling these eco-evolutionary processes. We leveraged 1,200 phages, 604 host genomes, and two decades of sampling at the same oyster farm to link ecological context with evolutionary outcomes across multiple biological scales. Four major insights emerged from our approach: (i) virulent phages can remain genetically stable and persist for years in an open marine environment; (ii) phage and bacterial populations coexist dynamically within oysters; (iii) MGEs drive genome plasticity, but their trajectories are shaped by clade structure and habitat-specific constraints; and (iv) oysters are hotspots of gene flow between vibrio, whose phylopathotypes harbor an unexpected diversity of MGEs, including novel episomal phage-plasmids and satellite-plasmids. Beyond their economic and ecological importance, oysters thus provide a natural laboratory where the interplay of ecology, evolution, and pathogenesis can be directly observed and experimentally dissected.

Our time-series revealed conservation of major vibrio clades and persistence of many virulent and temperate phages with identical genomes across four years. This stability is remarkable given the strong tidal currents at this Atlantic site, which should favor rapid microbial turnover. Importantly, the oysters used were hatchery-reared juveniles, newly introduced each season, and thus acted as “biological filters”, concentrating vibrio-phage assemblages from the surrounding seawater. Observed substitution rates (∼10⁻⁵ substitutions per site per year) indicate limited diversification. For temperate phages, persistence can be explained by vertical transmission as prophages, consistent with similarities between prophages detected in different years. Virulent phages pose a greater puzzle, as they are expected to persist only as particles; yet, we find they remain viable for years under laboratory storage (see Material and Method and Figure S31), suggesting similarly slow decay in the environment. Seasonal vibrio outbreaks then provide recurrent opportunities for infection. We propose that nested ecological filters explain this long-term stability: wild oysters sustain specific vibrio populations year-round^11^, those with appropriate colonisation factors. When conditions are inadequate for the propagation of the phage, the vibrio sustain temperate phages as prophages in their genomes and the virulent phage particles decay slowly enough to allow their propagation in the next season. The diversification of phages may then be constrained by environmental factors and receptor specificities. The key surprising observation is that despite the antagonistic co-evolution of phages and bacteria, via host defense and phage anti-defense systems, and that of phage and their parasite satellites, some phage lineages persist unaltered over several years. This suggests that evolutionary outcomes of phage-bacteria interactions in natural environments may be more constrained than usually acknowledged leading to rates of evolution markedly different from those reported in the lab.

A central aim of our study was to understand if and how phage and vibrio co-exist in oysters. We confirmed the clear ecological distinction between the environmental clade V1 and the oyster-associated phylopathotypes. Yet, we found that phylopathotype clades frequently coexist within the same animal, suggesting overlapping niches inside the oyster. We initially thought that phage pressure might regulate the type and abundance of vibrio over time and locations. A straightforward expectation of this hypothesis is that predator and prey abundances will be negatively correlated, as documented in some systems such as *V. cholerae* and its phage ICP1^26^. However, while both phage and bacterial abundances fluctuated over the four-month sampling interval, we saw no clear evidence that phage blooms were driving bacterial populations towards sharp decline. This may be caused by a diversity of mechanisms. Some vibrio strains may persist via phage resistance mechanisms or spatial refuges in oysters. Multiple phages can infect the same clade of vibrio, distributing predation pressure which results in lower correlations between individual phage and bacteria. Temperate phages may switch to lysogeny at high host densities^27^ reducing immediate lytic effects. Finally, other biotic and abiotic factors may impact vibrio population dynamics and satellites may impact the ones of phage.

Taken together, our results suggest that phages contribute to structuring vibrio populations, but not through simple cycles of dominance and collapse. Instead, within oysters, phage-host interactions are associated with long-term coexistence, with both antagonists persisting despite fluctuations in abundance and allowing recovery of the same phages and bacteria years apart.

The persistence of multiple bacterial and phage populations in oysters sets the stage for genetic exchanges driven by mobile genetic elements. We found that seawater-derived strains— particularly those from clade V1—have smaller genomes, lack plasmids, and remain consistently susceptible to phages. The relative scarcity of MGEs in these strains may reflect a trade-off that selects for reduced metabolic costs in the planktonic portion of the *V. crassostreae* population as noted in other surface bacterioplankton^28^. In contrast, oyster-associated strains carry larger genomes and significantly more MGEs, consistent with higher rates of gene turnover in this habitat. The oyster environment likely promotes MGE exchange through continuous nutrient inflow, extremely high microbial densities, and strong selective pressures from hemocyte immunity, antimicrobial peptides, metals^16,29,30^ and abundant phages (this study). Accordingly, we previously showed that genes involved in conjugative transfer and integrases are strongly upregulated during oyster colonization^31^, pointing to active MGE circulation *in vivo.* Interestingly, this ecological partitioning mirrors findings in other facultatively host-associated bacteria. For instance, freshwater *Escherichia coli* isolates tend to have smaller genomes and fewer MGEs than host-associated isolates, which carry more plasmids and phages^32^. The small genomes of *Pelagibacter ubique*, an extremely abundant marine species, have been streamlined to become completely devoid of mobile genetic elements^33^. Such parallels suggest that eukaryotic hosts act as hotspots of microbial genome plasticity that facilitate adaptation to both host and phage pressure, whereas aquatic habitats favor streamlined genomes.

We show that oysters are reservoirs of MGE diversity and hotspots for their exchange amongst vibrios. This high MGE diversity impacts adaptive processes because the types of elements differ in the traits they carry: plasmids tend to contribute to host adaptation and virulence, whereas prophages and satellites are enriched in genes for phage defense or inter-phage competition. Diversity can also result in novel modes of persistence. Over 6% of the temperate phages are phage-plasmids, combining the capacity for vertical inheritance as plasmids and long-range dispersal within viral particles. Similar elements have been described in marine *Vibrionaceae*, such as the *Autolykiviridae*^34^, which replicate as plasmids yet spread via the water column as virions. We also identified previously undescribed families of plasmid phage satellites (Cf-PISP and PISP), expanding the known diversity of MGEs in Bacteria. These novel elements provide the first examples of *Caudoviricetes* satellites that are maintained as plasmids and not as chromosomal islands. Unlike classical integrative satellites, these plasmid elements may exist in multiple copies per cell, which would facilitate the hijacking of helper phages for transmission between cells^35,36^. Similar elements have been detected through metagenomic sampling in the open ocean^35^ as particles containing plasmid-like replication genes (e.g. *parA, repA*), and in the recently described pDolos satellite-plasmid of *Shewanella*^36^. Satellite-plasmids can alter host evolution in distinctive ways—for example, by facilitating gene flow between satellite and plasmid gene pools^37^, or by maintaining higher effective copy numbers than chromosomally integrated elements which may increase mutation supply^38^. Our findings show that considering the ecology of marine bacteria is central to understanding the genetics of MGE biology in their ecosystem habitats.

Our study reveals that even in an open marine system, phage-host lineages can persist with little genomic change, stabilized by nested ecological filters rather than rapid turnover. Oysters provide a selective arena where microbial coexistence fosters both stability and innovation, concentrating mobile elements that shape bacterial adaptation and viral persistence. More broadly, host-associated environments that impose strong selective pressures and dense interactions may act as natural laboratories where mobile elements circulate and innovate, fueling the emergence of novel genetic forms such as phage–plasmids and satellites, while microbial evolution unfolds on ecological timescales.

## MATERIAL AND METHODS

### Sampling of *Vibrio* and phages in a marine environment

Samples were collected from an oyster farm in the Bay of Brest (Pointe du Château, 48° 20′ 06.19″ N, 4° 19′ 06.37″ W) three times per week from June 28 to September 15, 2021. Sampling began when seawater temperatures reached 16°C, a threshold associated with oyster mortalities^39^. Specific Pathogen-Free (SPF) juvenile oysters^40^ were deployed weekly (1,000 per batch) from June 1 to August 24, with mortality monitored at each sampling. When batch mortality exceeded 50%, subsequent sampling targeted oysters from the following week’s deployment. The study coincided with an ongoing oyster mortality outbreak lasting until September 9.

On each sampling date, 100 live oysters (≤50% mortality) were collected. Hemolymph was extracted, with individual samples (10 oysters) stored at -80°C for DNA extraction (Hi1–10; 35 dates = 350 DNA samples). The remaining hemolymph (90 oysters) was pooled, centrifuged, and filtered for viral analysis (35 dates = 35 samples). The pellet was resuspended in marine broth (MB), with fractions used for vibrio isolation and hemomicrobiota DNA storage (-80°C, 35 dates = 35 samples). Seawater (10 L) was size-fractionated as described previously^8^ with fractions resuspended and processed for DNA extraction (60, 5, 1, and 0.2 µm; 35 dates = 140 samples). Vibrios were isolated from 0.2 µm fractions, and microbiota samples were stored for further analysis (-80°C, 35 dates = 140 samples). Viral samples were concentrated 1,000-fold using iron chloride flocculation^41^, suspended in buffer, and stored at 4°C (viruses from seawater, 35 dates = 35 samples). Viral DNA was extracted from seawater and plasma samples (two sources, 35 dates = 70 DNA samples).

### Isolation and classification of *Vibrio crassostreae*

Total vibrio isolates from seawater (0.2 µm fraction) and hemolymph were cultured on Thiosulfate-Citrate-Bile Salts-Sucrose (TCBS) agar plates. From each sampling date, approximately 96 colonies were randomly selected from seawater and hemolymph plates and re-isolated once on TCBS. Putative *V. crassostreae* isolates were identified using multiplex PCR with three primer sets (Table S16), with colonies used as templates. Isolates were considered positive if at least two amplicons were obtained. To refine taxonomic identification, isolates were further analyzed by sequencing the *zrgB* gene, which discriminates among *V. crassostreae* phylogroups. Bacteria were grown overnight in marine broth (MB), and DNA was extracted using the Wizard extraction kit (Promega) according to the manufacturer’s instructions. The partial *zrgB* gene was amplified using conserved primers (Table S16), sequenced via Sanger sequencing (Eurofins), and used to construct a phylogenetic tree.

#### Phage isolation

To assess viral predation rates at the strain level, we used our previously established *Vibrio crassostreae* strain collection^9^, which includes 153 isolates from the Bay of Brest and Sylt (Germany), spanning clades V1 to V8. These strains served as bait for phage isolation across 35 sampling dates. For each date, we used a mix of 10 µL of seawater viral concentrate (1,000×, equivalent to 10 mL of seawater) and 10 µL of plasma derived from a pooled sample of 90 oysters. Phage infections were detected by plaque formation using soft agar overlays on bacterial lawns. In total, 5,355 interactions were tested (35 dates × 153 strains).

To refine strain-level predation rate estimates across spatial and temporal dimensions, we focused on host-date combinations that initially exhibited more than 20 Plaque Forming Units (PFU) per plate, with hosts belonging to a specific clade. These combinations were further tested using either 10 µL of seawater viral concentrate or 10 µL of plasma.

In a previous study^9^, we collected up to six plaques per morphotype and found that they were mostly clonal. Therefore, we purified one phage per plaque morphotype and combination (host and date), resulting in a final collection of 1331 phages. Phages were re-isolated through up to three rounds of plaque purification to ensure purity. High-titer stocks (>10⁹ PFU/mL) were prepared via confluent lysis in agar overlays and stored at 4°C, with an additional aliquot stored at -80°C in the presence of 25% glycerol. After four years, isolates remained viable, with titers reduced on average by ∼1 log (Figure S31).

#### Primer design and ddPCR protocol

To identify SNP-dense regions specific to each *V. crassostreae* subpopulation, we used Snippy (https://github.com/tseemann/snippy) for variant calling, using GV1681 as the reference genome. SNPs were mapped across 604 *V. crassostreae* genomes, including the 447 isolates from this study, with sensitivity and specificity calculated for each subpopulation. Only SNPs in core genes with optimal sensitivity and specificity were retained. SNPs were then treated as nodes in a graph, with edges weighted by proximity, and filtered based on primer design criteria (25–40 bp primer length, 180–300 bp amplicon size, and at least two specific positions per primer). Protein multiple sequence alignments (mafft) were converted to CDS alignments, and SNPs were mapped onto these using GV1681 as a reference. Final primer and probe selection was validated through visual inspection. For phages, primer design was simplified by selecting core genes unique to the subpopulation of interest but absent in other phages.

Droplet digital PCR reactions were performed either in single plex reactions using an intercalant DNA dye or in multiplex using dye-labeled specific probes. For single target amplification, reactions consisted of 20 μL mixture per well containing 10 μL of ddPCR Evagreen Supermix, 600 nM of primers (Table S16) and 5 μL of DNA. For multiple target amplifications, reactions consisted of 20µL per well containing 5 µL of ddPCR Multiplex Supermix, 900nM of primers, 250nM of dye-labeled specific probes (Table S16) with either FAM, HEX, Cy5 or Cy5.5, 4mM of DDT and 5uL of DNA. The ddPCR reactions were incorporated into droplets using the QX100 Droplet Generator (Bio-Rad). Nucleic acids were amplified with the following cycling conditions: 5 min at 95 °C, 40 cycles of 30 s at 95 °C and 60 °C for 60 s using a Bio-Rad’s C1000 Touch Thermal Cycler. For the single plex reactions, an extra final droplet cure step of 5 min at 4°C then 5 min at 90 °C was incorporated. Droplets were read and analyzed using Bio-Rad QX600 system and QuantaSoft software (version 1.7.4.0917) in ‘absolute quantification’ mode. Only wells containing ≥10,000 droplets were accepted for further analysis.

### *Vibrio crassostreae* genome sequencing and assembly

Of 512 confirmed *V. crassostreae* isolates, 453 (89%) clustered into distinct clades and were selected for genome sequencing, while 59 isolates (11%) were more diverse and excluded from this study. A total of 447 vibrios were successfully sequenced at the Biomics platform of the Pasteur Institute using Illumina® TruSeq™ DNA PCR-Free High Library Prep Kit, IDT for Illumina-TruSeq DNA UD Indexes. Runs were performed by 96 samples with a first validation on MiSeq flow cell micro v2, paired-end 2×150 cycles, next a production run on NovaSeq 1 lane, paired-end 2×150 cycles.

To improve the genome assembly, 20 *V. crassostreae* strains from our previously established *V. crassostreae* strain collection^9^, distributed across the phylogeny, were sequenced by PacBio. The PacBio genome assembly was conducted with Unicycler (version 0.4.9)^42^, leveraging a hybrid approach that combines both Illumina short reads and PacBio long reads. Post-assembly processing involved Python scripts for replicon identification, where contigs are classified into chromosomes and plasmids, terminal contig sequence extension using raw Illumina reads to ensure completeness and searching the assembled contigs against the closest relative genomes. The final manual curation step involved identifying and correcting RNA stretches within the assembled contigs, conducting manual alignments to further refine the assembly and ensure its accuracy, resolving any ambiguous bases that may have emerged during the assembly process, and circularizing the contigs to accurately reflect the complete genomic structure.

*Vibrio* isolates without Pacbio hybrid sequencing data were de-novo assembled using Illumina short reads with post-assembly scaffolding as follows. Reads were trimmed using Trimmomatic v0.39 (LEADING:3, TRAILING:3, SLIDINGWINDOW:4:15, MINLEN:36)^43^. *De novo* read assembly was performed using Spades version 3.15.2 (--careful --cov-cutoff auto -k 21,33,55,77 -m 10). Contig circularization was manually confirmed by inspection of the assembly graph with bandage v0.9.0^44^. To improve syntenic consistency, the contigs of each strain were scaffolded along the completed assembly of the most closely related of the 20 strains sequenced with PacBio using RagTag v2.1.0 scaffold^45^ with minimap2 v2.24 r1122^46^.

#### Phage genome sequencing and assembly

The methodology for phage DNA extraction, sequencing, and classification followed the protocol detailed previously^47^. Briefly, phage high titer stocks were concentrated using PEG precipitation, treated with nucleases, and subjected to phenol-chloroform extraction. DNA integrity was assessed via agarose gel electrophoresis and quantified using Qubit. Phage sequencing was performed at the Biomics platform of the Pasteur Institute (Paris, France). DNA was fragmented using Covaris (target size: 500 bp), and libraries were prepared with the TruSeq™ DNA PCR-Free High Library Prep Kit. Due to inefficiency in adaptor ligation, an amplification step of 14 cycles was added using Illumina P7 and P5 primers (IDT). Sequencing was carried out on a MiSeq Micro v2 flow cell with paired-end 2×150 cycles.

Phage isolate reads were trimmed using Trimmomatic v0.39^43^ and assembled *de-novo* with SPAdes v3.15.2. Contaminant contigs were filtered using the UniVec Database, and the resulting one-contig phage genome was manually linearized.

#### Genome annotation of bacteria, plasmids, phages, and satellites

Genes were predicted and annotated in *V. crassostreae* genomes using bakta v1.9.2^48^ with full database v5.1. Except where otherwise indicated, tools which require predicted coding sequences were provided with the corresponding amino acid or nucleotide sequences predicted by bakta.

Annotations in 5 representative high-quality plasmid sequences were manually curated and refined. Conjugative systems and Mob relaxases were predicted and classified using CONJscan v2.0.1 and MOBscan respectively with default parameters^49,50^.

Sequenced lytic phages, predicted prophages, and phage-satellites were annotated using pharokka v1.7.3^51^ using default settings and database v1.4.0. Automatic annotations were subsequently manually refined in certain prophages and phage-satellites. HMMsearch v3.4^52^ and the HMM profiles packaged with SatelliteFinder v0.9^53^ were used to identify satellite-associated genes. Genes of unknown function were classified by comparing translated amino acid sequences against online databases using the blastp web portal^54^ to screen against non-redundant proteins in RefSeq release 229 with default alignment parameters and using the InterProScan web portal to screen against all available protein signature databases^55^. Structural comparisons were conducted by predicting the structure of unknown proteins as monomeric proteins using AlphaFold3 Server to screen against the available structural databases of FoldSeek in 3Di/AA mode^56,57^.

### Inference of plasmid presence and transfer in *V. crassostreae*

Twenty plasmids identified in the 20 Pacbio hybrid-assembled genomes were manually classified by inspection of gene content into five plasmid families: pGV, p1, pAlioth, pMintaka, and pMizar. To identify the relationships within each family, core genes were identified as MMseqs2 v15-6f452^58^ protein clusters with sensitivity 7.5, minimum coverage 80%, coverage-mode 5, and minimum sequence ID 80%. Core gene nucleotide sequences were aligned with MAFFT v7.526^59^--globalpair and the unpartitioned concatenated alignment was used to construct a phylogeny for each plasmid using iqtree v2.3.6 with MFP identifying GTR+F+I as the best model by Bayesian information criterion^60,61^.

Illumina reads from each sequenced *V. crassostreae* strain were cleaned with fastp v0.24.0 and remapped with minimap2 v2.28 r1209 with flag-x sr to each of the 20 plasmid sequences to assemble hypothetical plasmids of each family. Protein coding regions were predicted from the samtools v1.20 consensus^62^ remapped sequences with prodigal v2.6.3^63^ using the –p meta procedure, as recommended for plasmids. Weighted gene repertoire relatedness (wGRR) was calculated as previously described^37^ between each plasmid and the reference used during remapping and a plasmid marked as present in a strain if wGRR was greater than 50%. To infer transfer between strains, the plasmid in each group with the highest wGRR was marked as the most closely related plasmid.

Cleaned reads were used to relative extrachromosomal element copy numbers by remapping, as described above, to the corresponding genome assemblies. Read depths were calculated with samtools v1.20 coverage. Read depths were normalized to the mean chromosomal read depth in each strain.

#### *In silico* prediction of prophages, phage-satellites

Prophages were predicted with geNomad v1.8.1 end-to-end^17^ using database version 1.7 and default settings. The prophages predicted by geNomad as non-chromosomally integrated (direct terminal repeats or no terminal repeats) were manually inspected for circularization in *de novo* assemblies. As a filter for genome quality, prophages were only retained for analysis if they were predicted *Caudoviricetes* with a genome size greater than 25 kbp, predicted *Tectiliviricetes* with a genome size of greater than 10 kbp, or predicted *Faserviricetes* with a genome size of greater than 4 kbp, in accordance with the literature^18^.

Phage satellites were detected using SatelliteFinder v0.9^53^ containerized in Apptainer^64^ with MacSyFinder v2.0^65^ using the predicted protein sequence output of bakta with flags --linear and --ordered_replicon. Predictions were made with each of the four available satellite model definitions bundled with the software: PICI, cf-PICI, PLE, and P4.

#### Distribution of genes between mobile genetic elements

Regions of genomic plasticity (RGPs) were identified using the panRGP module^66^ of ppanggolin version 2.2.1^67^ to identify the persistent and flexible regions of the pangenome in the full corpus of 605 *V. crassostreae* strains. This identified 32,349 RGPs which were then classified into their MGE types. RGPs were classified as plasmids (3217 RGPs) if they were identified by geNomad as having likely plasmid origin. RGPs were classified as satellites (450 RGPs) if they contained an entire predicted phage satellite segment predicted by SatelliteFinder. RGPs were classified as prophages (4478 RGPs) if they were either identified by geNomad as containing a prophage sequence or by checkV v1.0.3^68^ with database release 1.5 as a complete, low, medium, or high-quality prophage containing at least one viral gene, as previously described^9^. Finally, the presence of integrase genes was predicted in the RGPs (1945 RGPs) using bakta, as above, or annotated as unclassified (22259 RGPs) if none of the above applied.

Anti-phage defense genes were inferred using padloc v2.0.0^69^ with prodigal v2.6.3 for gene prediction and hmmer v3.3.2 to screen against database release 2.0.0. Biocide and metal resistance genes were identified by screening for the genes included in the experimental confirmed gene database of BacMet2^70^ with MMseqs with minimum sequence identity 70% and minimum coverage 50%. Antimicrobial resistance genes were identified by screening for the genes included in the protein homolog CARD database v4.0.1^71,72^ with MMseqs with minimum sequence identity 30% and minimum coverage 50%. Virulence factors were identified by screening for the genes included in the VFDB core dataset^73^ with MMseqs with minimum sequence identity 70% and minimum coverage 50%. An in-house Python script was used to check for the localization of these systems within the flexible genome regions predicted by panRGP. Comparisons of coding density were performed for the predictions of each dataset using a Kruskal-Walis H-test with Benjamini-Hochberg correction for false discovery rate and with Dunn’s post-hoc test implemented in scipy and scikit_posthocs respectively.

### *Vibrio crassostreae* core-genome alignment and phylogenetic inference

Bacterial persistent genome phylogeny was conducted using the PanACoTA workflow^74^ version 1.3.1-dev2, without applying Mash v2.3 distance filtering^75^. Prodigal version v2.6.3^63^ was used for gene prediction. Persistent gene families were thresholded as being present in 90% or greater of all genomes after MMseqs clustering on an 80% protein identity threshold. IQ-TREE v2.0.3^61^ was used for phylogenetic tree inference using the core-genome concatenation, applying the GTR model with 1000 bootstrap replicates. The total pangenome comprised 29,988 families, while the persistent genome contained 3,099 families.

The gene presence-absence table and phylogenetic tree from the PanACoTA workflow were used to perform a phylogeny-aware pangenome analysis using panstripe v0.3.1^76^ with ‘”family = gaussian” parameter provided to model fit. Comparisons of pangenome characteristics between subsets of the *V. crassostreae* population were conducted using pruned subtrees of the same base phylogenetic tree to maintain a comparable scale.

Phylogenetic comparative methods which leveraged tree topology for correction of statistical non-independence were performed using the phylolm v2.6.5^77^ and phytools v2.4-4^78^ packages of R v4.4.1. Specifically, continuous data was analyzed using phylANOVA with 10000 iterations for p-value simulation with Holm-Bonferroni post-hoc testing. The association between genome size and the number of MGEs detected was tested with phylolm using a Brownian motion tree model. Presence/absence of MGEs was modeled as a binary trait using phyloglm with a logistic MLE regression.

#### Phage and prophage genomic and functional classification

The entire lytic phage collection (including both new and previously-collected isolates) was clustered using the Viral Intergenomic Distance Calculator (VIRIDIC)^13^. VIRIDIC (v1.1) generates a single output file for all blastn results, which can be a limiting factor when dealing with large datasets. To address this, a custom script was developed to split the blastn outputs into smaller, manageable files, allowing for better parallelization and more efficient processing of the 1,268 phage genomes. VIRIDIC utilizes fastcluster for fast hierarchical clustering of the resulting distance matrix. The taxonomic assignment of viruses into family, genus, and species ranks was performed based on the intergenomic similarity thresholds defined by the International Committee on Taxonomy of Viruses (ICTV): genus (≥70% similarity), and species (≥95% similarity). Additionally, lysogeny scores were computed for each lytic phage using BACPHLIP v0.9.6^79^ using default options. Clustering of intergenomic sequence identity of predicted prophages was performed using VIRIDIC in the same way. Prophages were taxonomically assigned following the ICTV’s Viral Metadata Resource release 19 based on marker gene taxonomy during geNomad prediction.

Visualizations of the *V. crassostreae* clades used to isolate lytic phages were produced using the R package ggraph 2.2.1^80^ to draw hierarchical connections between phage genera, phage species, and phage isolates. Isolates are colored according to the clade of the *Vibrio* host used as bait during isolation. A similar network shows the link between *Caudoviricetes* prophage genera, species, and individual predicted prophages, while the color of the leaf prophage shows the clade of the *Vibrio* genome from which the prophage was predicted.

#### Inference of substitution rates in persistent lytic and temperate phage species

We considered a lytic phage species to be persistent if it was isolated in both the 2017 and 2021 sampling campaigns. Similarly, we considered a temperate phage species to be persistent if its lysogens were collected in two or more different years. We generated whole-genome alignments between phage genomes of a single phage species using mafft --globalpair. Because the PHI test implemented in PhiPack v1.0^81^ inferred evidence for recombination in LS32, Gubbins v3.4^82^ was used to mask recombinant regions with default settings for tree construction, model fitting, and sequence reconstruction^83–85^. Final tree inference and inference of substitution rate was performed using BEAST v2.6.3^86^ with 100,000,000 iterations and 10% burn-in and a coalescent constant population size model with an exponential relaxed clock rate and a clock rate prior of 1.9 x 10^-4^ site^-1^year^-1^ (the estimated recombination-free substitution rate in a population of lactophages^15^). Path sampling implemented in the model-selection BEAST package^87,88^ did not support strict clock rate or Bayesian skyline population size^89,90^ as alternative model specifications.

#### Satellite-plasmid detection and comparison

SatelliteFinder was used to characterise TG54, a 15 kbp contig identified by geNomad as a putative *Caudoviricetes*. Though SatelliteFinder did not identify TG93 as a satellite, inspection of its annotated gene content lead us to identify it as a cfPICI-like element. TG54 (cf-PISP) encodes a PFam family PF05065 capsid protein and TG93 (PISP) encodes a PFam family PF03864 CapsidPE protein. These HMM profiles were downloaded from the InterPro repository^91,92^. HMMsearch v3.4 [http://hmmer.org/] was used with default parameters to identify homologous proteins of each capsid family in these new elements, in the phage-satellite sequences included with the supplementary data of^53^ and in the lytic vibriophages catalogued in this manuscript. For each capsid family, gene sequences were aligned with mafft –linsi and a maximum-likelihood tree was inferred using IQtree with the Q.pfam+F+I+R model specified. The most similar phage-satellite to each of our novel elements was chosen using these capsid phylogenies and used, together with the prototypical PICI or cfPICI with the most experimental characterization^24,25,93^ for comparisons of genetic organisation. Comparisons were produced by computing bi-directional best hits with MMseqs and visualized with pygenomeviz v1.4.1 https://github.com/moshi4/pyGenomeViz.

We looked for similar elements in public repositories by screening for similarities to the capsid or capsid_PE and the parA protein sequences to identify other species carrying plasmid-satellites. We searched for similar sequences using the blastp web portal in non-redundant proteins in RefSeq release 229 with default alignment parameters. From the top 100 hits for each search, the genetic contexts of the target protein were manually inspected for similarity to the plasmid- satellite gene catalogue and for the size of the contig and its integration into the host chromosome. Each homolog was downloaded and re-annotated with pharokka v1.7.3 using phanotate for gene prediction with the expectation that coding density and gene length distribution in these elements is best modeled as phage-like. Syntenies were produced as above.

## ACKNOWLEDGEMENTS

We thank Bruno Petton (Ifremer Brest) for providing oyster juveniles and overseeing in situ experiments, which made the time-series sampling possible. We are grateful to Chloé Berger, Pauline Daszkowski, Justine Groseille, Théo Foutel Rodier, Étienne Levêque, and Mariam Mamba (GV team, Station Biologique de Roscoff, France) for their assistance during sampling and collections and Carine Diarra (Le Roux lab, Montréal) for technical help. We thank Jorge Moura de Sousa for helpful discussions concerning the biology and classification of phage-satellites and Julien Guglielmini for help running wGRR/GRIS. We thank Léa Verena Zinsli and Otto X. Cordero for valuable comments on the manuscript, and Illumina for reduced pricing of sequencing kits. Editorial support from Stephen Matheson (Life Science Editors) is also warmly acknowledged. This research was enabled in part by the computational resources of Calcul Quebec and the Digital Research Alliance of Canada.

This work was supported by funding from the European Research Council (ERC) under the European Union’s Horizon 2020 research and innovation program (grant agreement No. 884988, Advanced ERC Dynamic), the Canada Excellence Research Chairs Program (CERC-2022-00051), and the Fonds de recherche du Québec – Nature et technologies (FRQ-NT, Fonds des leaders John-R.-Evans, grant 44584) awarded to FLR. Substantial support was provided by the Agence Nationale de la Recherche (ANR-20-CE35-0014 “RESISTE”) and the Fonds UdeM-Pasteur pour la découverte de nouveaux antibiotiques et antibactériens, a philanthropic fund at the Courtois Institute in Biomedical Innovation, Faculty of Medicine, University of Montreal to EPCR and FLR. Biomics Platform, C2RT, Institut Pasteur, Paris, France, is supported by France Génomique (ANR-10-INBS-09) and IBISA.

## AUTHOR CONTRIBUTIONS

FLR conceived the study, FLR and EPCR supervised the project and secured funding. KC, DP, DCG, YL, LM and FLR conducted the experiments. JL, DG, CB and EPCR performed the genomic analyses. JL, KC, DP, DCG, EPCR and FLR analysed the data. JL, EPCR and FLR, wrote the manuscript.

## COMPETINGS INTERESTS

Authors declare no competing interests.

## DATA AVAILABILITY STATEMENT

### Data availability

The sequenced genomes have been deposited in the NCBI database, with accession numbers listed in Supplementary Table S1 and S2.

### Materials availability

The phage and their vibrio hosts have been deposited in the following public collection: https://roscoff-culture-collection.org/ with accession numbers listed in Tables S1 and S3.

