## Supplementary Figures for "Ecological constraints foster both extreme viral-host lineage stability and mobile element diversity in a marine community"

### SUPPLEMENTARY MATERIAL

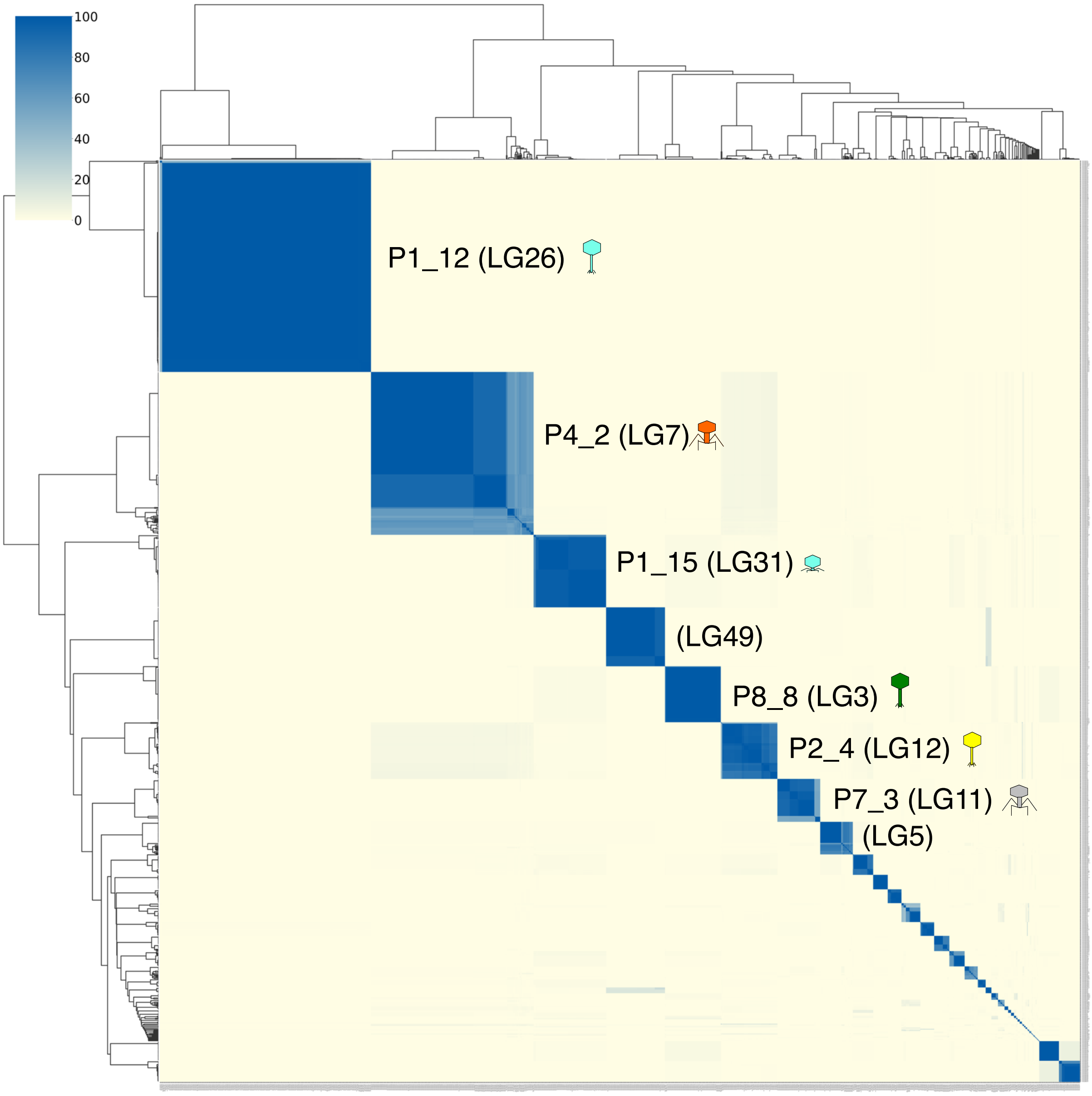


**Figure S1** Phages isolated in the present study were taxonomically classified using VIRIDIC. The most prominently represented clusters are denoted by their respective genus names, as per the classification outlined by Piel et al., 2022. Genus numbers assigned in this study are shown in brackets. When known, the corresponding morphotype is illustrated with an icon.

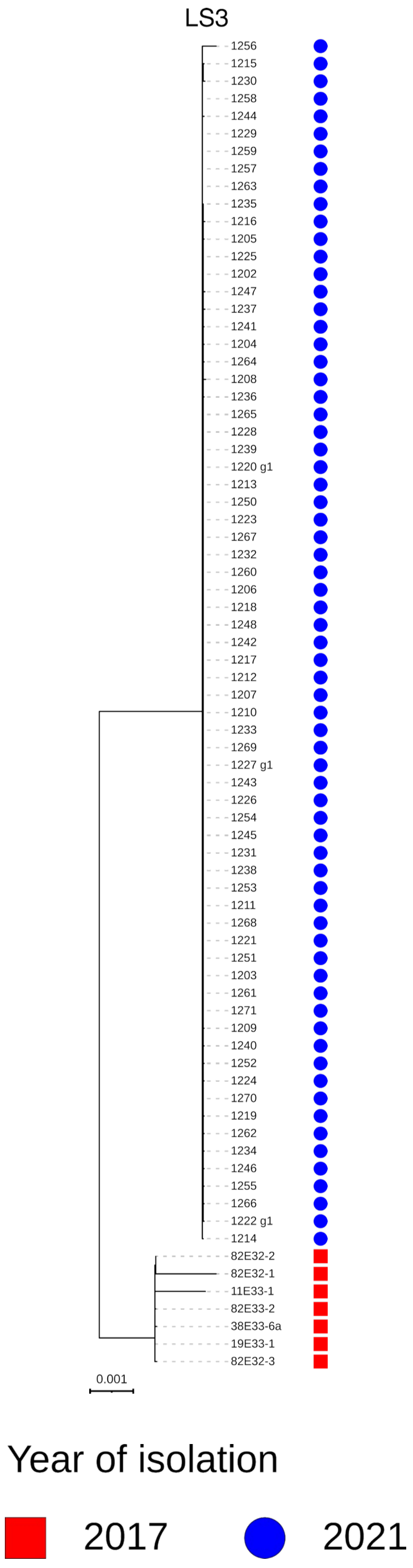


**Figure S2 The inferred phylogeny of virulent phages from species LS3 isolated in both 2017 and 2021.** Maximum-likelihood trees were generated from whole genome alignments of sequenced phage genomes and are displayed as midpoint-rooted. Annotations to the right indicate the year of isolation for each phage. Annotations follow the convention of Figure 2 with red squares showing isolates from 2017 and blue circles showing isolates from 2021.

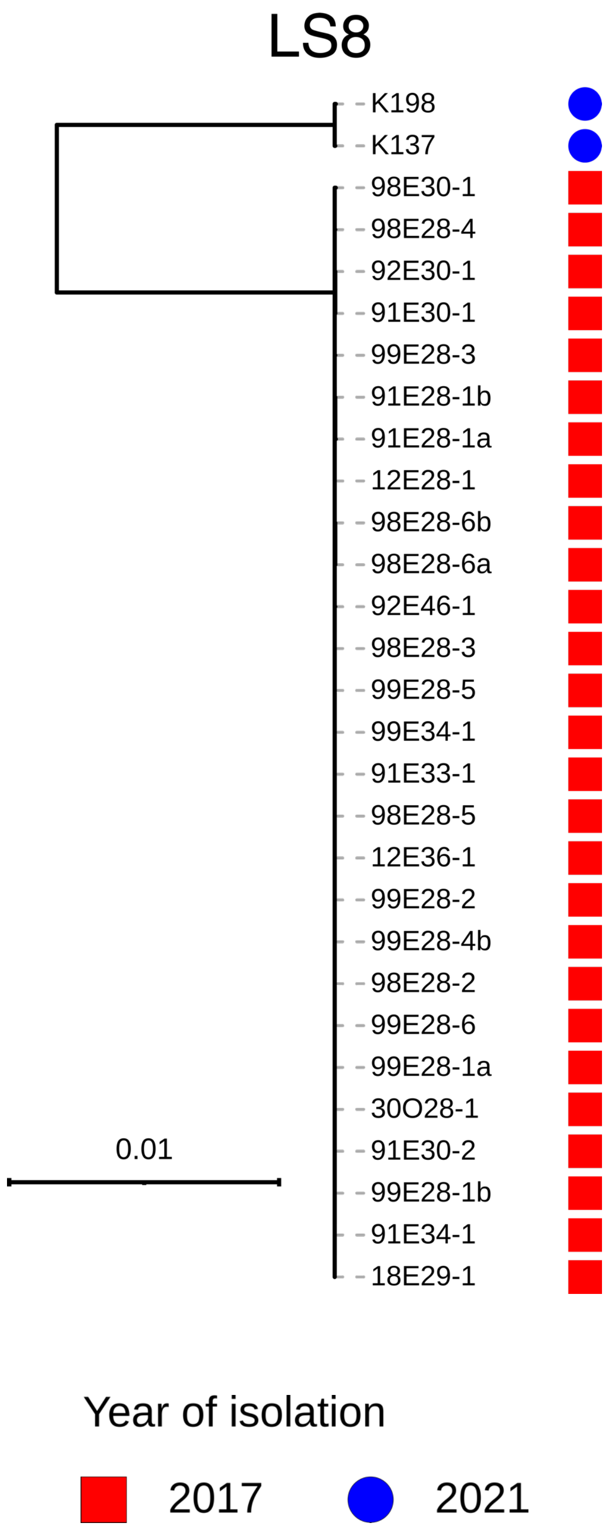


**Figure S3 The inferred phylogeny of virulent phages from species LS8 isolated in both 2017 and 2021.** Maximum-likelihood trees generated from whole genome alignments of sequenced phage genomes are displayed as midpoint-rooted. Annotations to the right indicate the year of isolation for each phage. Annotations follow the convention of Figure 2 with red squares showing isolates from 2017 and blue circles showing isolates from 2021.

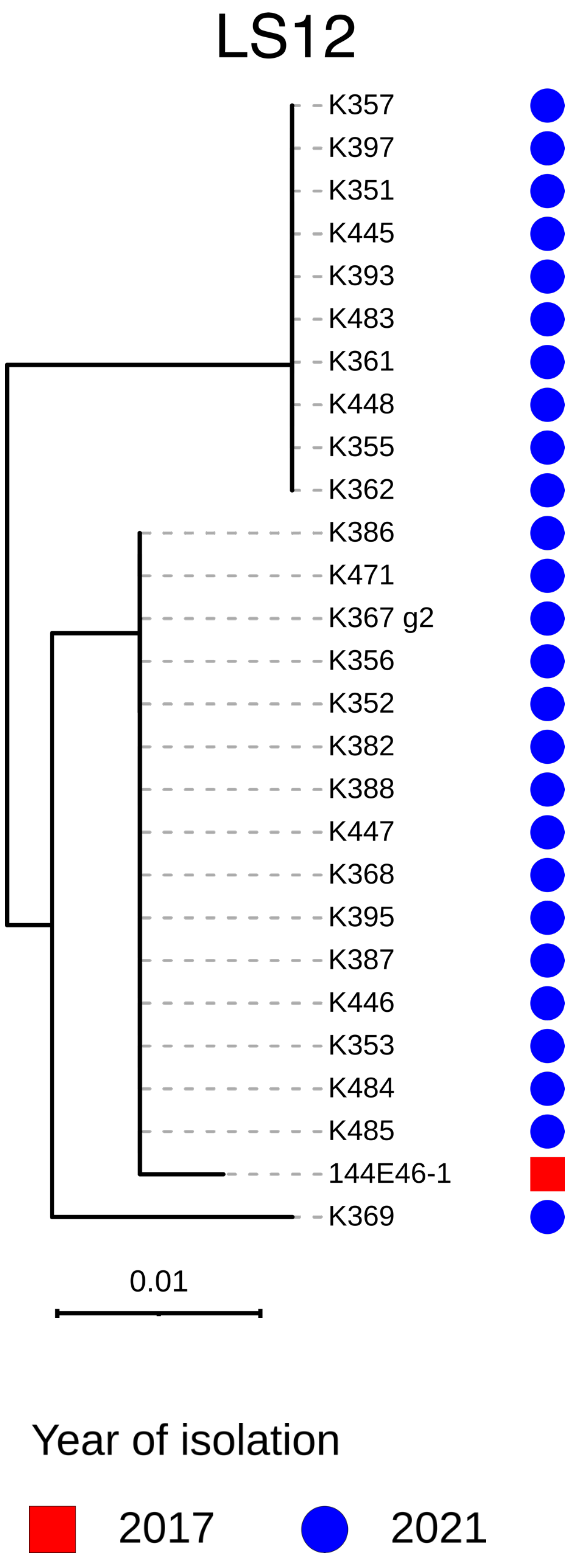


**Figure S4 The inferred phylogeny of virulent phages from species LS12 isolated in both 2017 and 2021.** Maximum-likelihood trees generated from whole genome alignments of sequenced phage genomes are displayed as midpoint-rooted. Annotations to the right indicate the year of isolation for each phage. Annotations follow the convention of Figure 2 with red squares showing isolates from 2017 and blue circles showing isolates from 2021.

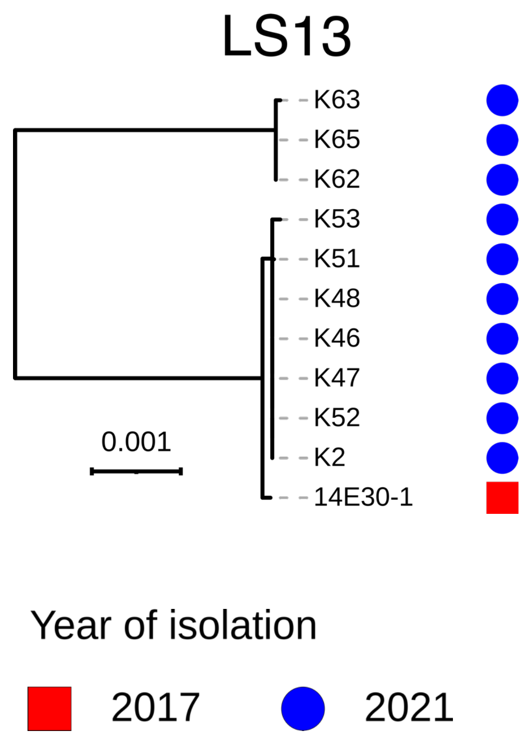


**Figure S5 The inferred phylogeny of virulent phages from species LS13 isolated in both 2017 and 2021.** Maximum-likelihood trees generated from whole genome alignments of sequenced phage genomes are displayed as midpoint-rooted. Annotations to the right indicate the year of isolation for each phage. Annotations follow the convention of Figure 2 with red squares showing isolates from 2017 and blue circles showing isolates from 2021.

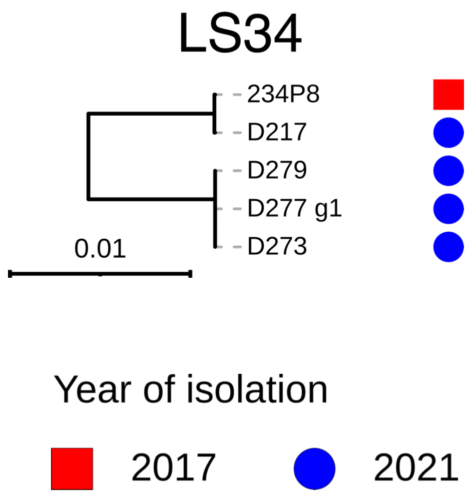


**Figure S6 The inferred phylogeny of virulent phages from species LS34 isolated in both 2017 and 2021.** Maximum-likelihood trees generated from whole genome alignments of sequenced phage genomes are displayed as midpoint-rooted. Annotations to the right indicate the year of isolation for each phage. Annotations follow the convention of Figure 2 with red squares showing isolates from 2017 and blue circles showing isolates from 2021.

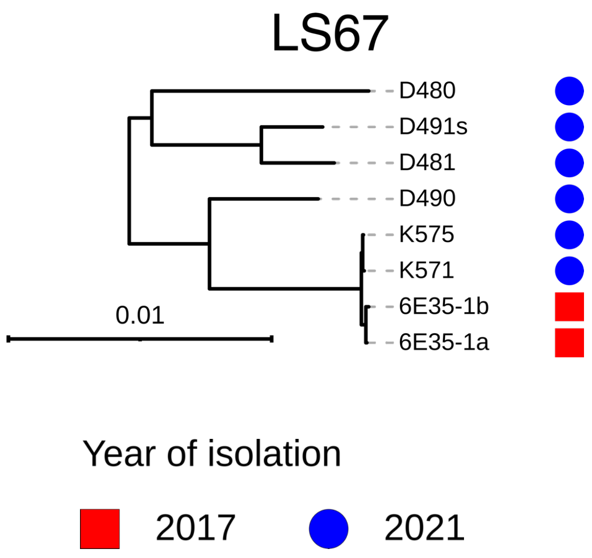


**Figure S7 The inferred phylogeny of virulent phages from species LS67 isolated in both 2017 and 2021.** Maximum-likelihood trees generated from whole genome alignments of sequenced phage genomes are displayed as midpoint-rooted. Annotations to the right indicate the year of isolation for each phage. Annotations follow the convention of Figure 2 with red squares showing isolates from 2017 and blue circles showing isolates from 2021.

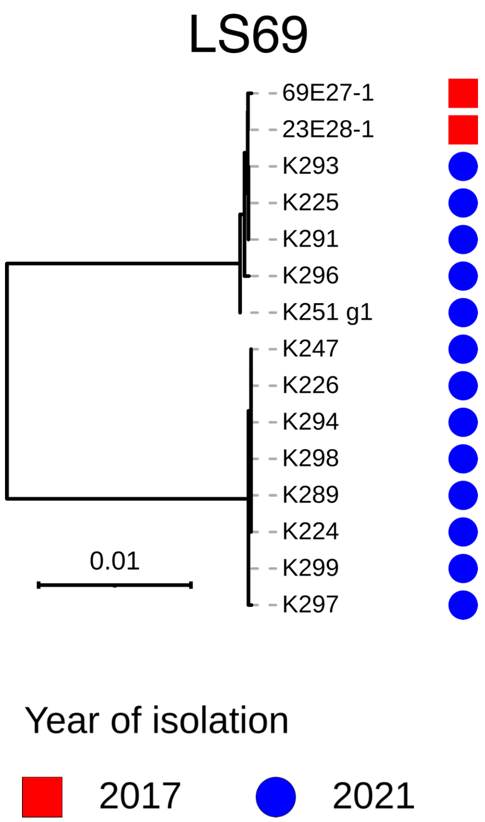


**Figure S8 The inferred phylogeny of virulent phages from species LS69 isolated in both 2017 and 2021.** Maximum-likelihood trees generated from whole genome alignments of sequenced phage genomes are displayed as midpoint-rooted. Annotations to the right indicate the year of isolation for each phage. Annotations follow the convention of Figure 2 with red squares showing isolates from 2017 and blue circles showing isolates from 2021.

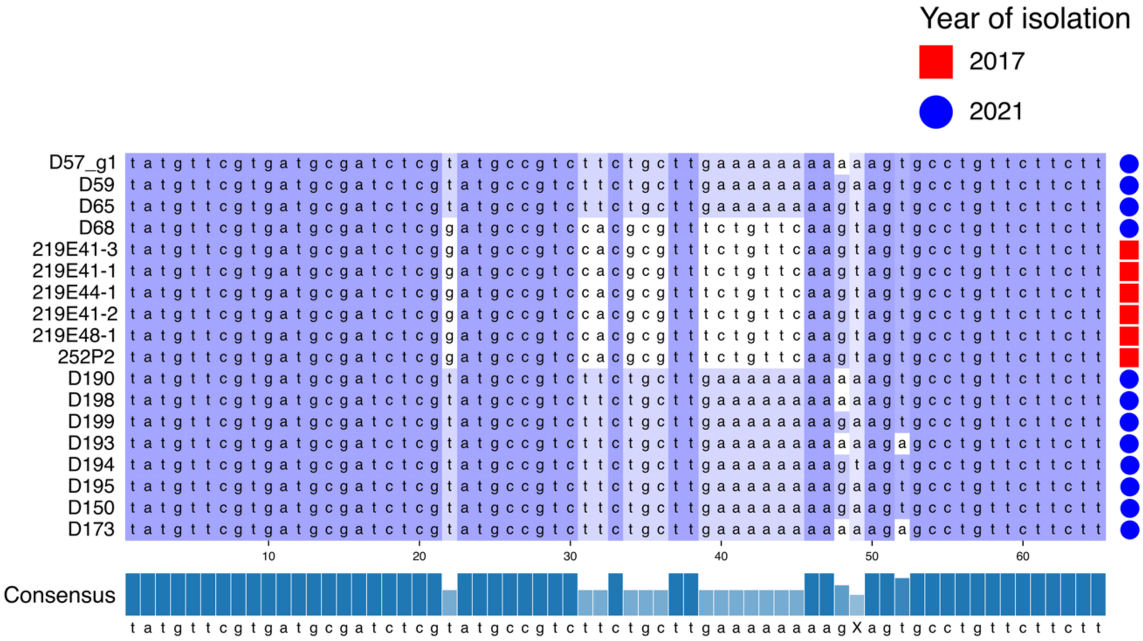


**Figure S9 Intergenomic variation in LS29 is largely localized to a short region within a gene of unknown function.** A section of a whole genome alignment between 18 phages of LS29 shows 16 variable sites which are a major component of the phylogenetic distance separating 2021 phage isolates from 2017 phages (and D68). The year of isolation of each phage is indicated to the right, showing that D68 preserves the ancestral sequence. Sequences are colored by percent conservation at each site and sequence consensus is shown in the bottom bar.

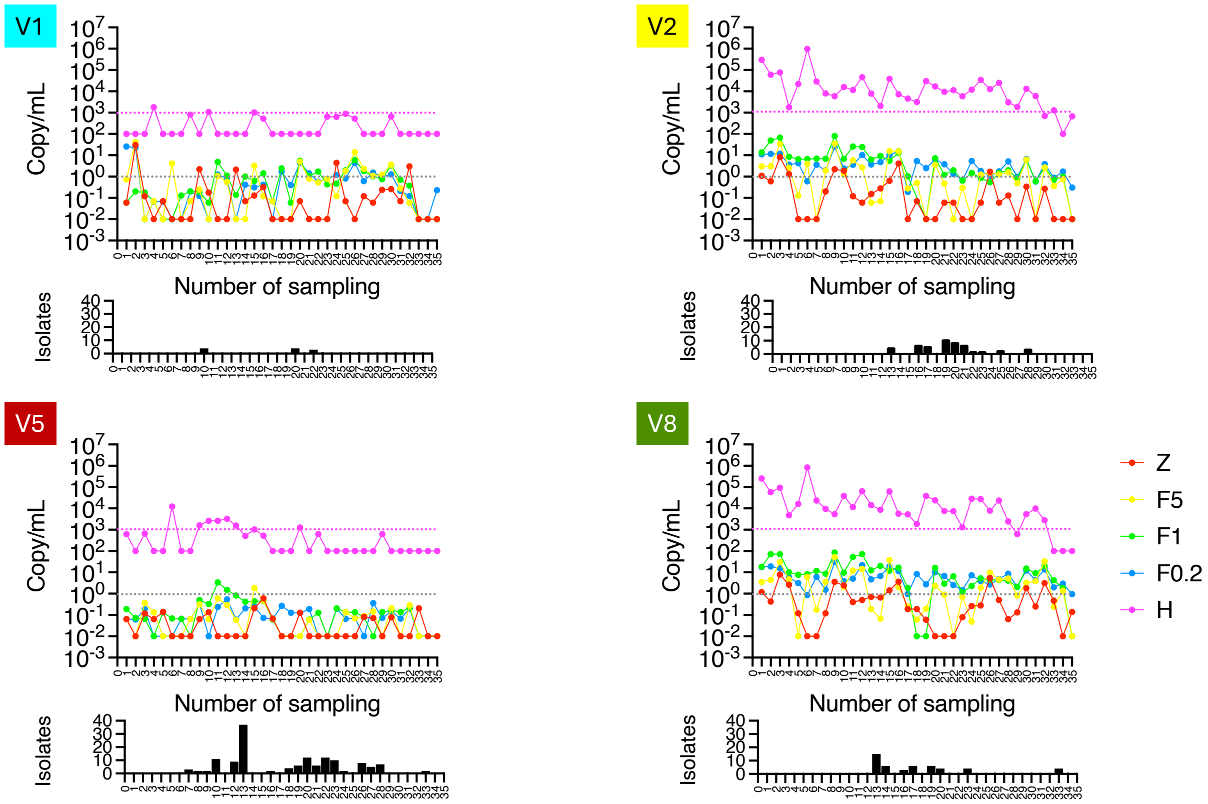


**Figure S10 Quantification of *V. crassostreae* clades in seawater and oyster hemolymph by digital droplet PCR (ddPCR).** Seawater was size-fractionated (Z>60 µm; F5: 60-5 µm; F1: 5-1 µm; F0.2: 1-0.2 µm; H: hemolymph) and DNA extracted from each fraction, while hemolymph from 90 oysters was pooled for DNA extraction. Each point represents the absolute DNA copy number per mL of hemolymph or seawater. The dotted horizontal line indicates the detection limit. Histograms below each graph show the number of strains isolated per date.

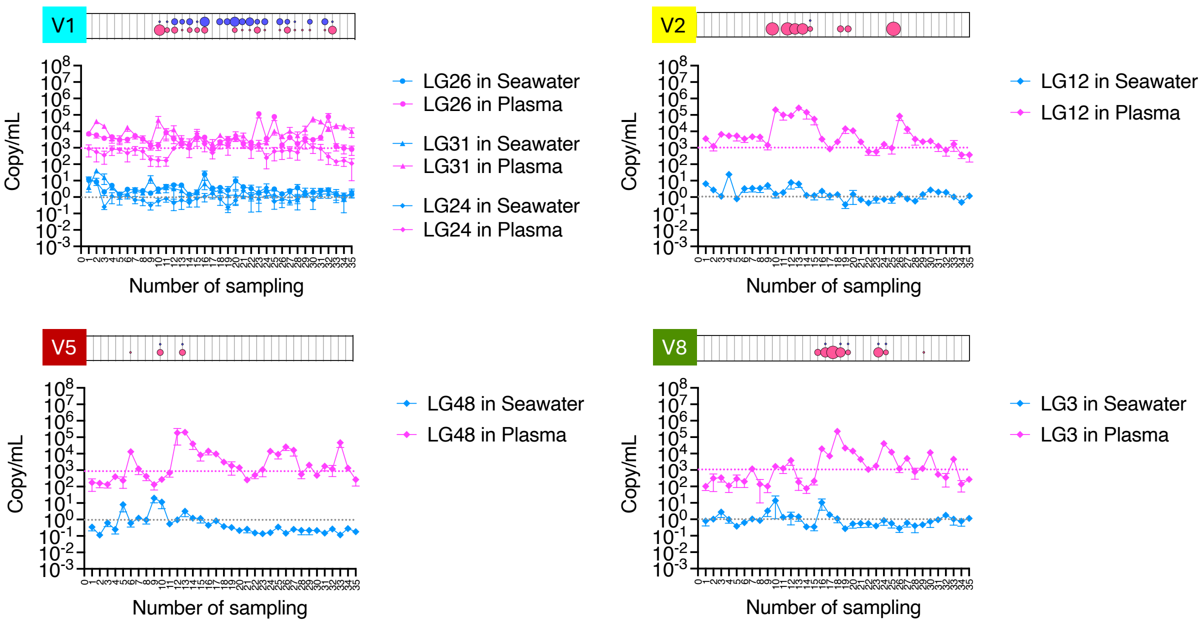


**Figure S11 Quantification of phage genera in seawater and oyster plasma by digital droplet PCR (ddPCR).** Viral fractions from seawater (<0.2 µm) were concentrated 1,000-fold using iron chloride flocculation, while viral fractions from plasma (<0.2 µm) were obtained from pooled hemolymph of 90 oysters. Viral DNA was extracted and quantified by ddPCR. Each point represents the absolute DNA copy number per mL of plasma or seawater. The dotted horizontal lines indicate the detection limit. Dot plots above each graph show average PFU counts per clade and sampling date, measured from the same samples (10 µL of seawater concentrate, equivalent to 10 mL of raw seawater, or 10 µL of oyster plasma; see Figure 3A), providing an independent validation of the ddPCR data.

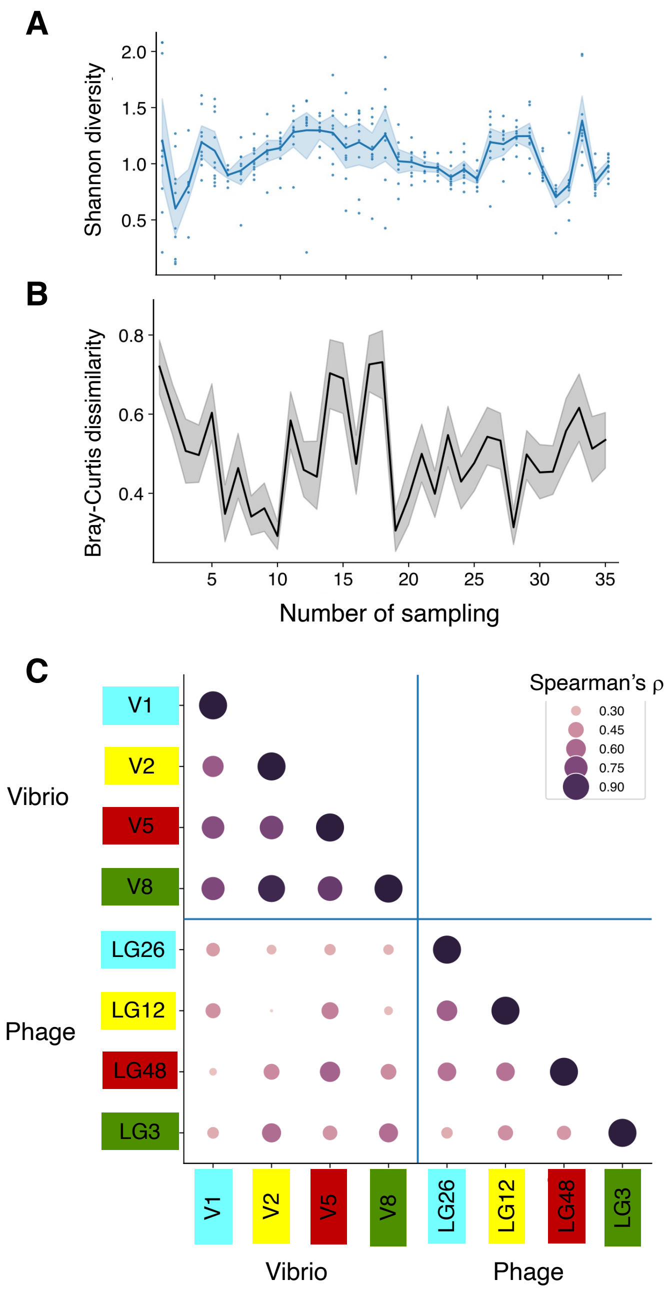


**Figure S12 Diversity indices of oysters infected by Vibrio clades and their corresponding lytic phage genera (as shown in Figure 2), based on ddPCR-determined DNA copy number per mL. A** Alpha-diversity, measured using the Shannon index, shows temporal fluctuations over the sampling period but no clear overall trend. Each point represents an individual oyster; the solid line indicates the mean, with shading denoting the 95% confidence interval. **B** Beta-diversity**,** calculated as the mean Bray–Curtis dissimilarity across all 45 pairwise comparisons of 10 oysters per date, reflects variability in the composition of vibrio clades or phage genera. Periods of low beta diversity (i.e., more uniform composition across oysters) often coincide with population blooms. Shading shows 95% confidence interval. **C** Spearman correlations between tracked populations reveal strong co-fluctuations among vibrio clades but surprisingly weak correlations between each vibrio clade and its corresponding phage genus.

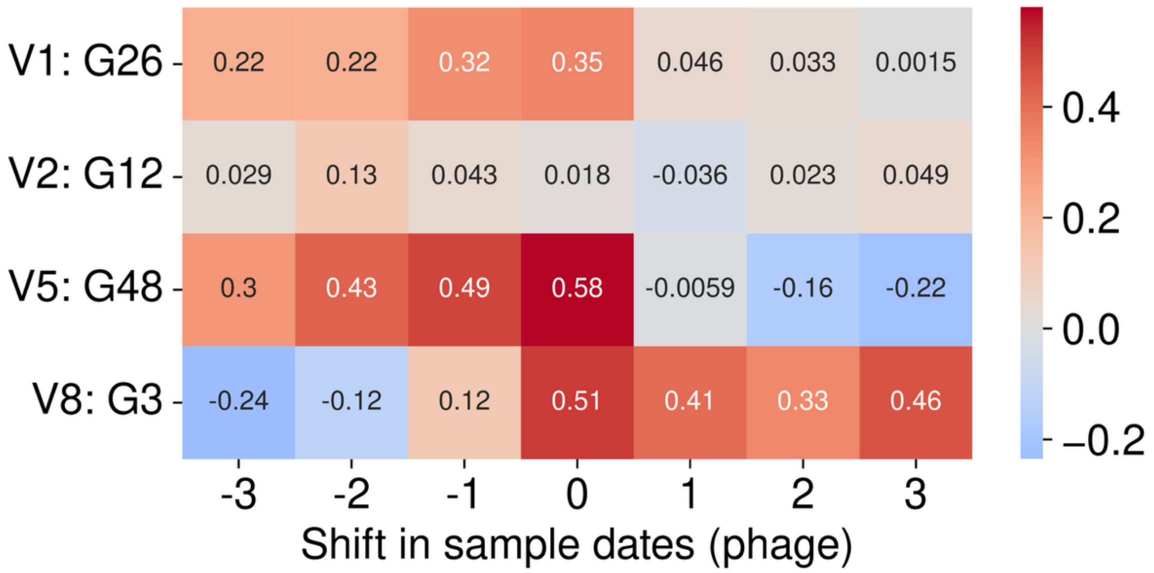


**Figure S13 Nucleic acid–based quantification of *V. crassostreae* populations and their co-infecting phages in oyster hemolymph shows no strong correlations.** Vibrio and phage abundances were quantified in hemolymph from ten individual oysters across 35 sampling dates using ddPCR with lineage-specific primers. For each sampling date, Spearman's rho was calculated between the geometric mean of V. crassostreae clade abundances and the corresponding phage genus. Correlations were assessed for contemporaneous samples (lag = 0) as well as with the phage series shifted relative to Vibrio (negative = earlier phage, positive = later phage) to test for temporal lags in phage replication on susceptible hosts. No correlations were significant (Table S2).

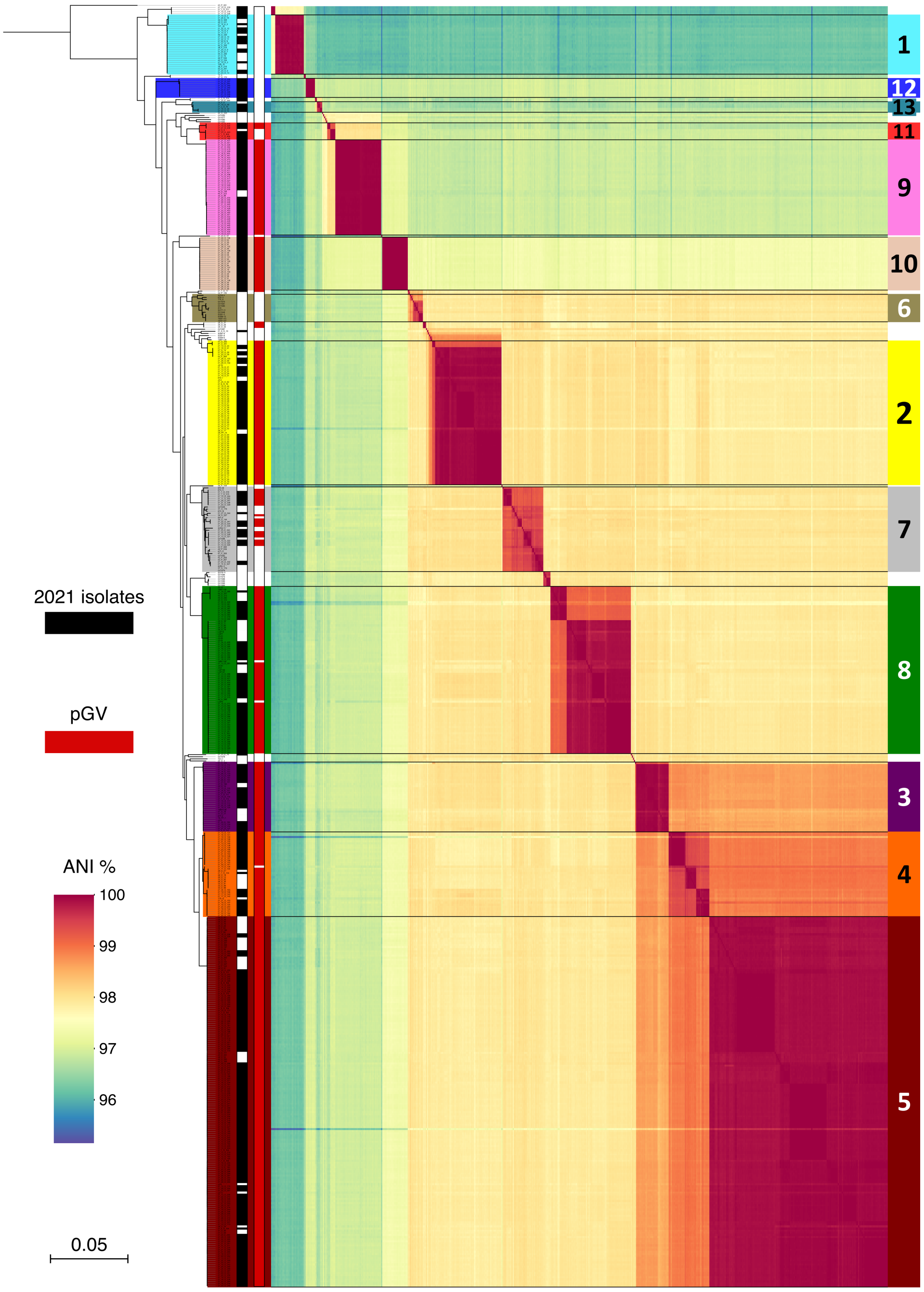


**Figure S14 Categorization of bacterial isolates into clades within the *V. crassostreae* species.** Core genome phylogeny based on 3,099 gene families of 605 *V. crassostreae* isolates, with pairwise Acid Nucleic Identities (ANI) values revealing highly organized clades within the species. Clade designations and corresponding colors Refer to Piel et al., 2022 with new clades designated V9 to V13. The second column represents sampling, with 2021 dates in black and other collections in white. The presence of the virulence-associated plasmid, pGV, is highlighted in red in the second column.

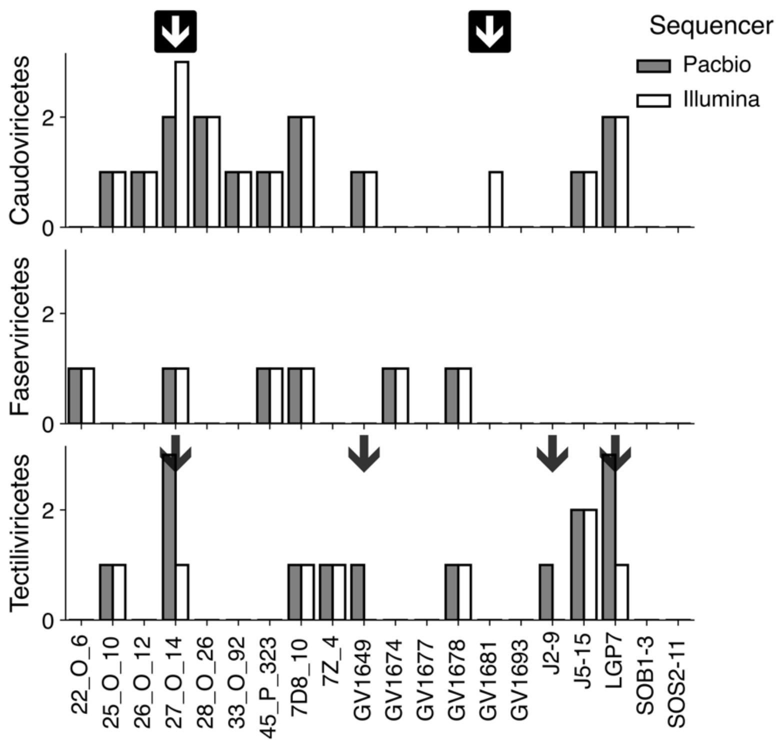


**Figure S15 Comparison of prophage detection in 20 *V. crassostreae* genomes sequenced with both PacBio and Illumina platforms.** Gaps in the identification of temperate *Tectiliviricetes* phages are indicated by arrows**.**

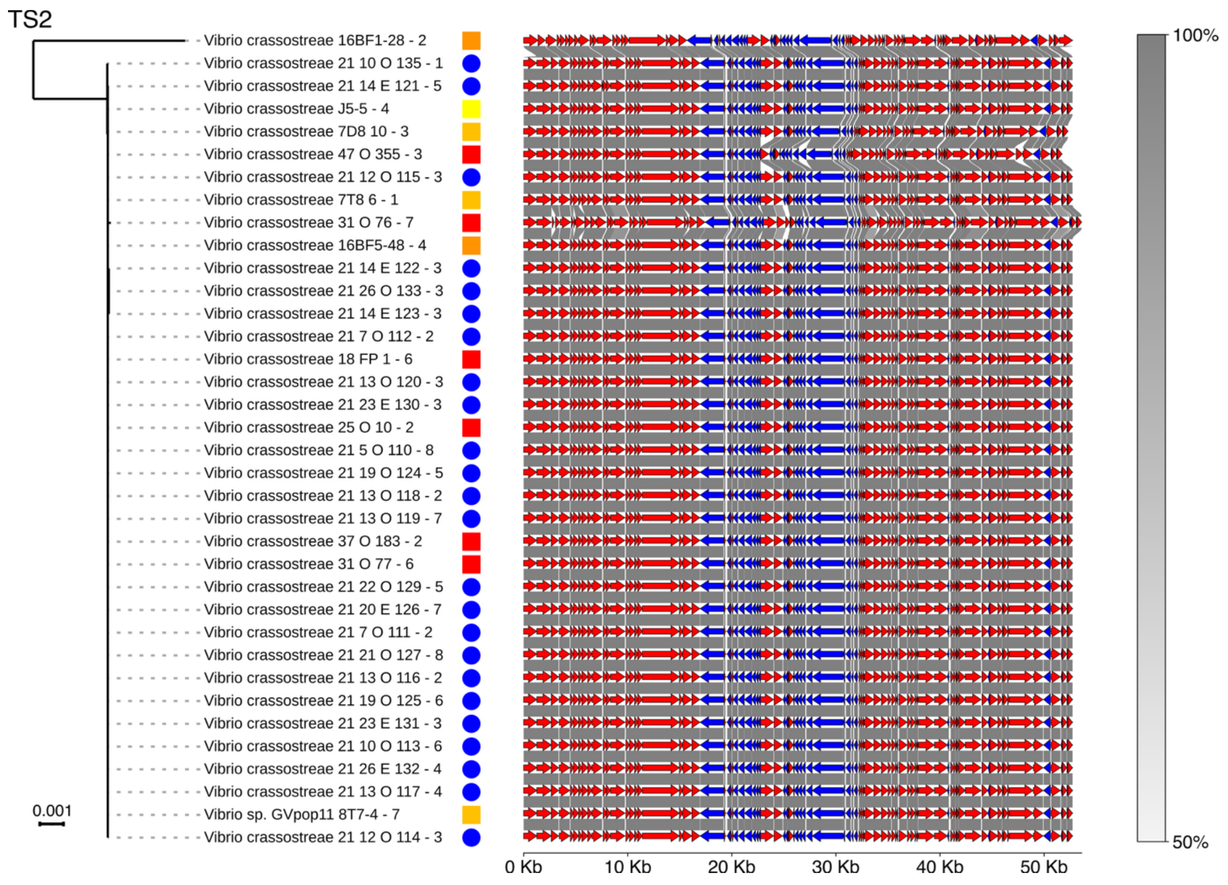


**Figure S16 Phylogeny of temperate phage species TS2 identified in *V. crassostreae* strains isolated between 2001 and 2021.** Maximum-likelihood trees generated from whole-genome alignments of phage genomes are displayed as midpoint-rooted. Annotations to the right indicate the year of isolation for each phage, following the color code shown in the legend. Visualization of synteny shows each temperate phage genome with red and blue arrows showing sense and anti-sense ORFs predicted using pharokka. Links between genes in adjacent genomes show similarity between bidirectional best hits calculated using MMseqs2 and visualized using pygenomeviz.

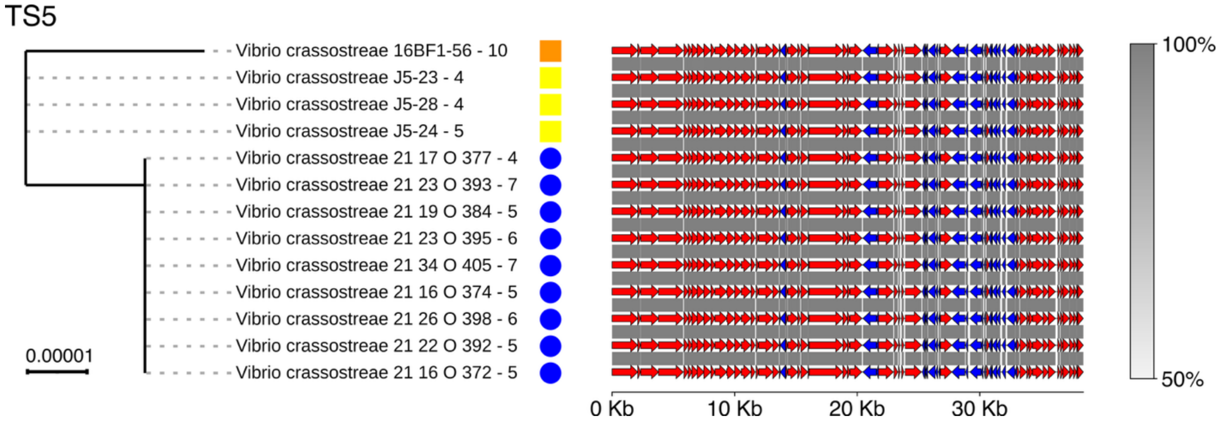


**Figure S17 Phylogeny of temperate phage species TS5 identified in *V. crassostreae* strains isolated between 2001 and 2021.** Maximum-likelihood trees generated from whole-genome alignments of phage genomes are displayed as midpoint-rooted. Annotations to the right indicate the year of isolation for each phage, following the color code shown in the legend. Visualization of synteny shows each temperate phage genome with red and blue arrows showing sense and anti-sense ORFs predicted using pharokka. Links between genes in adjacent genomes show similarity between bidirectional best hits calculated using MMseqs2 and visualized using pygenomeviz.

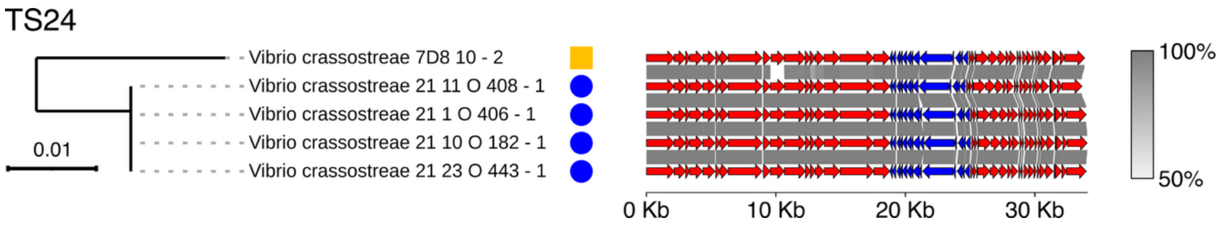


**Figure S18 Phylogeny of temperate phage species TS24 identified in *V. crassostreae* strains isolated between 2001 and 2021.** Maximum-likelihood trees generated from whole-genome alignments of phage genomes are displayed as midpoint-rooted. Annotations to the right indicate the year of isolation for each phage, following the color code shown in the legend. Visualization of synteny shows each temperate phage genome with red and blue arrows showing sense and anti-sense ORFs predicted using pharokka. Links between genes in adjacent genomes show similarity between bidirectional best hits calculated using MMseqs2 and visualized using pygenomeviz.

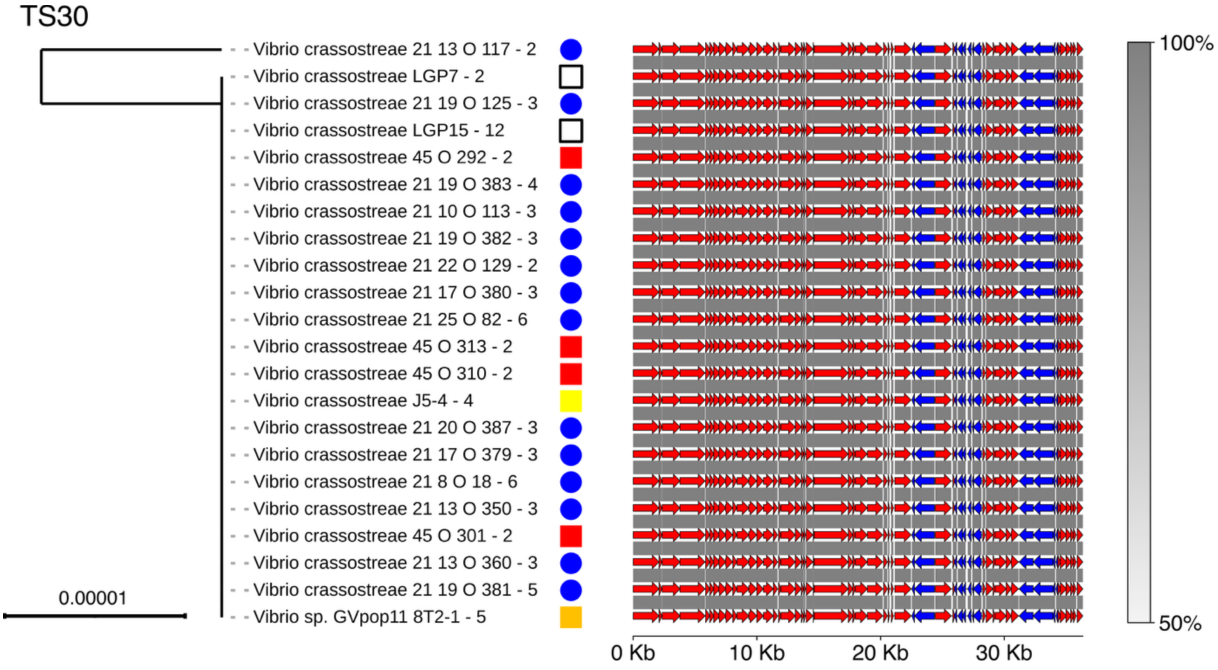


**Figure S19 Phylogeny of temperate phage species TS30 identified in *V. crassostreae* strains isolated between 2001 and 2021.** Maximum-likelihood trees generated from whole-genome alignments of phage genomes are displayed as midpoint-rooted. Annotations to the right indicate the year of isolation for each phage, following the color code shown in the legend. Visualization of synteny shows each temperate phage genome with red and blue arrows showing sense and anti-sense ORFs predicted using pharokka. Links between genes in adjacent genomes show similarity between bidirectional best hits calculated using MMseqs2 and visualized using pygenomeviz.

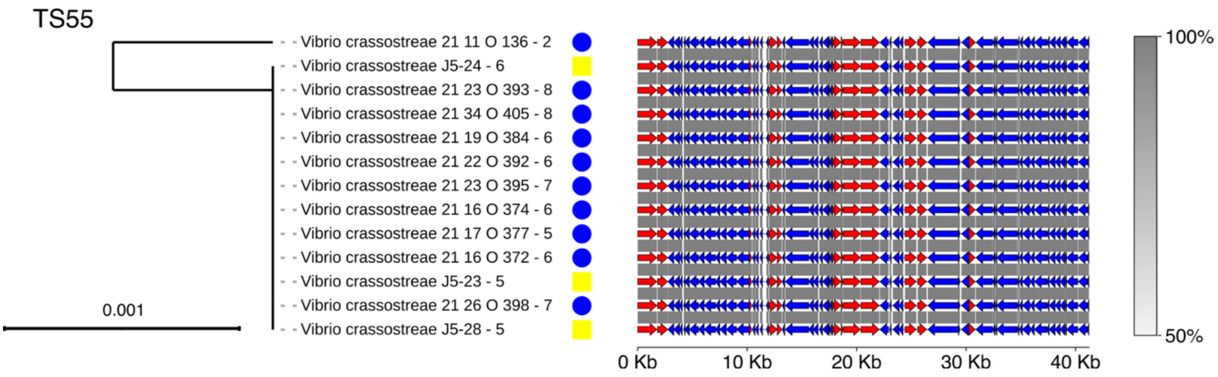


**Figure S20 Phylogeny of temperate phage species TS55 identified in *V. crassostreae* strains isolated between 2001 and 2021.** Maximum-likelihood trees generated from whole-genome alignments of phage genomes are displayed as midpoint-rooted. Annotations to the right indicate the year of isolation for each phage, following the color code shown in the legend. Visualization of synteny shows each temperate phage genome with red and blue arrows showing sense and anti-sense ORFs predicted using pharokka. Links between genes in adjacent genomes show similarity between bidirectional best hits calculated using MMseqs2 and visualized using pygenomeviz.

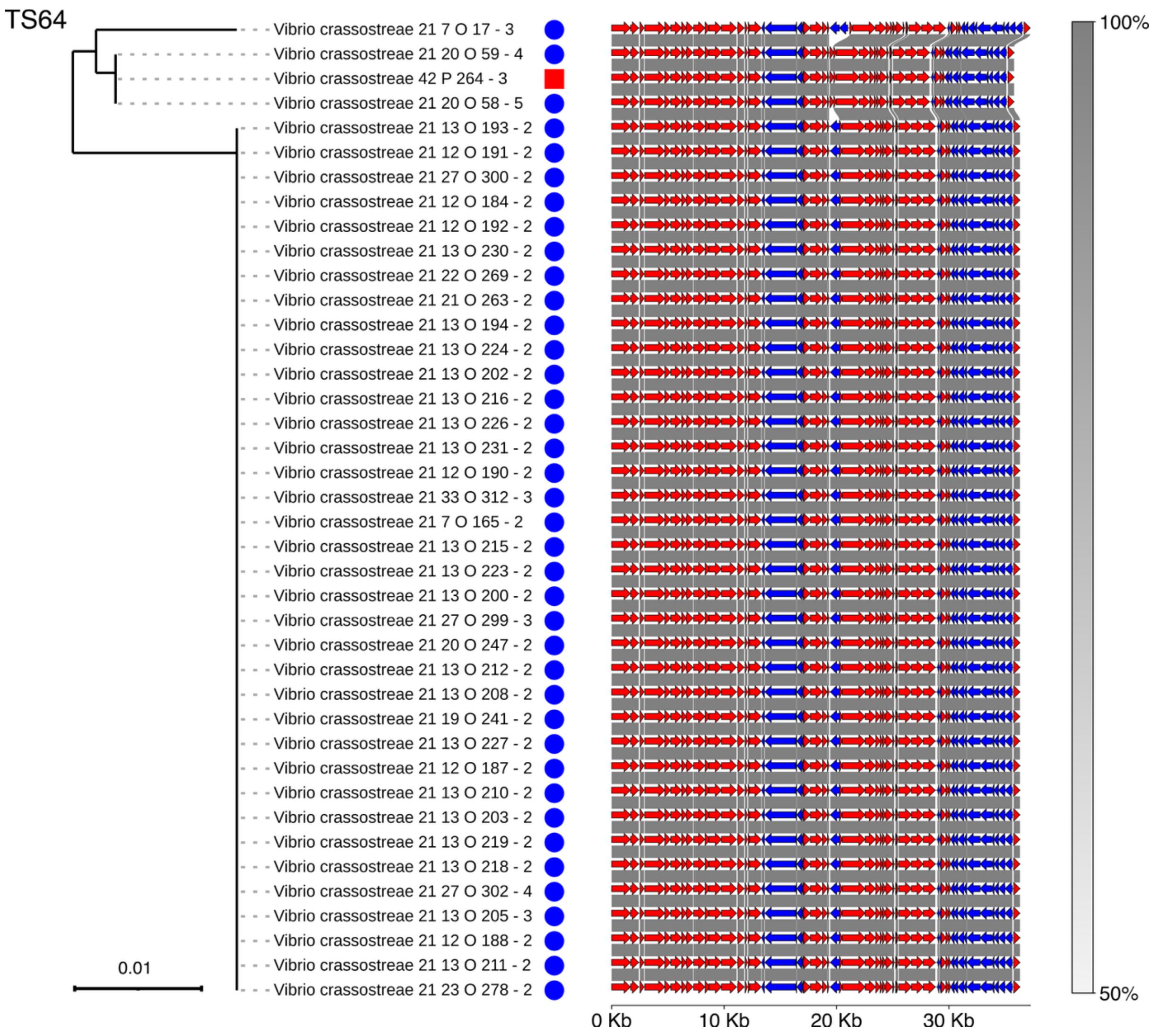


**Figure S21 Phylogeny of temperate phage species TS64 identified in *V. crassostreae* strains isolated between 2001 and 2021.** Maximum-likelihood trees generated from whole-genome alignments of phage genomes are displayed as midpoint-rooted. Annotations to the right indicate the year of isolation for each phage, following the color code shown in the legend. Visualization of synteny shows each temperate phage genome with red and blue arrows showing sense and anti-sense ORFs predicted using pharokka. Links between genes in adjacent genomes show similarity between bidirectional best hits calculated using MMseqs2 and visualized using pygenomeviz.

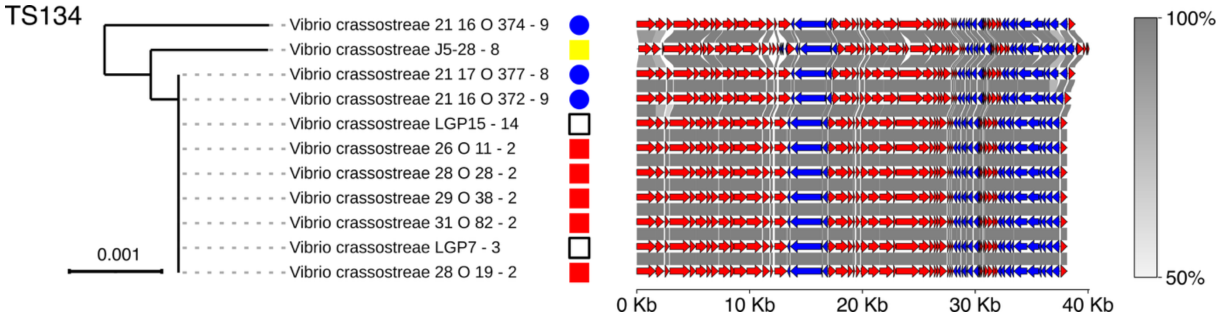


**Figure S22 Phylogeny of temperate phage species TS134 identified in *V. crassostreae* strains isolated between 2001 and 2021.** Maximum-likelihood trees generated from whole-genome alignments of phage genomes are displayed as midpoint-rooted. Annotations to the right indicate the year of isolation for each phage, following the color code shown in the legend. Visualization of synteny shows each temperate phage genome with red and blue arrows showing sense and anti-sense ORFs predicted using pharokka. Links between genes in adjacent genomes show similarity between bidirectional best hits calculated using MMseqs2 and visualized using pygenomeviz.

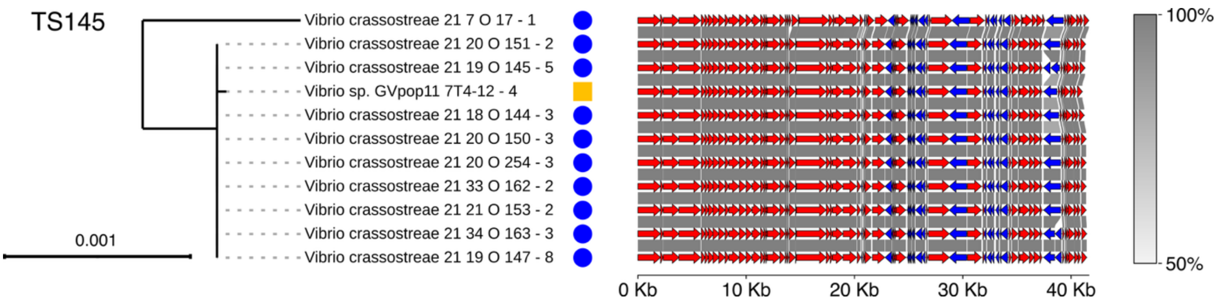


**Figure S23 Phylogeny of temperate phage species TS145 identified in *V. crassostreae* strains isolated between 2001 and 2021.** Maximum-likelihood trees generated from whole-genome alignments of phage genomes are displayed as midpoint-rooted. Annotations to the right indicate the year of isolation for each phage, following the color code shown in the legend. Visualization of synteny shows each temperate phage genome with red and blue arrows showing sense and anti-sense ORFs predicted using pharokka. Links between genes in adjacent genomes show similarity between bidirectional best hits calculated using MMseqs2 and visualized using pygenomeviz.

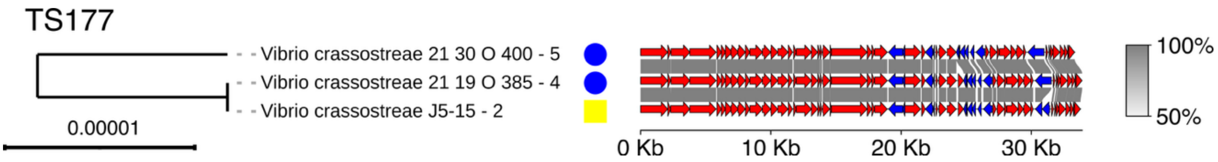


**Figure S24 Phylogeny of temperate phage species TS177 identified in *V. crassostreae* strains isolated between 2001 and 2021.** Maximum-likelihood trees generated from whole-genome alignments of phage genomes are displayed as midpoint-rooted. Annotations to the right indicate the year of isolation for each phage, following the color code shown in the legend. Visualization of synteny shows each temperate phage genome with red and blue arrows showing sense and anti-sense ORFs predicted using pharokka. Links between genes in adjacent genomes show similarity between bidirectional best hits calculated using MMseqs2 and visualized using pygenomeviz.

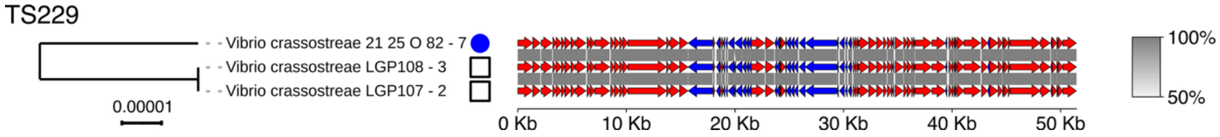


**Figure S25 Phylogeny of temperate phage species TS229 identified in *V. crassostreae* strains isolated between 2001 and 2021.** Maximum-likelihood trees generated from whole-genome alignments of phage genomes are displayed as midpoint-rooted. Annotations to the right indicate the year of isolation for each phage, following the color code shown in the legend. Visualization of synteny shows each temperate phage genome with red and blue arrows showing sense and anti-sense ORFs predicted using pharokka. Links between genes in adjacent genomes show similarity between bidirectional best hits calculated using MMseqs2 and visualized using pygenomeviz.

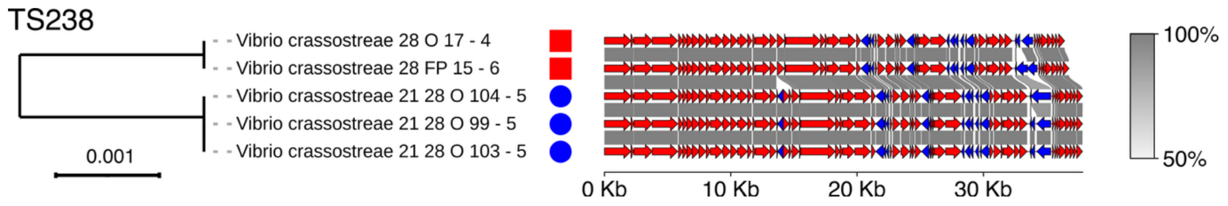


**Figure S26 Phylogeny of temperate phage species TS238 identified in *V. crassostreae* strains isolated between 2001 and 2021.** Maximum-likelihood trees generated from whole-genome alignments of phage genomes are displayed as midpoint-rooted. Annotations to the right indicate the year of isolation for each phage, following the color code shown in the legend. Visualization of synteny shows each temperate phage genome with red and blue arrows showing sense and anti-sense ORFs predicted using pharokka. Links between genes in adjacent genomes show similarity between bidirectional best hits calculated using MMseqs2 and visualized using pygenomeviz.

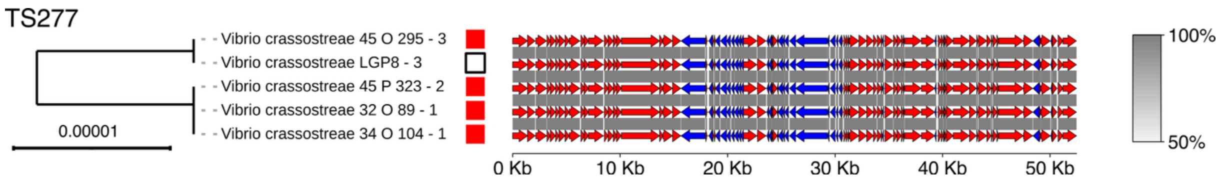


**Figure S27 Phylogeny of temperate phage species TS277 identified in *V. crassostreae* strains isolated between 2001 and 2021.** Maximum-likelihood trees generated from whole-genome alignments of phage genomes are displayed as midpoint-rooted. Annotations to the right indicate the year of isolation for each phage, following the color code shown in the legend.

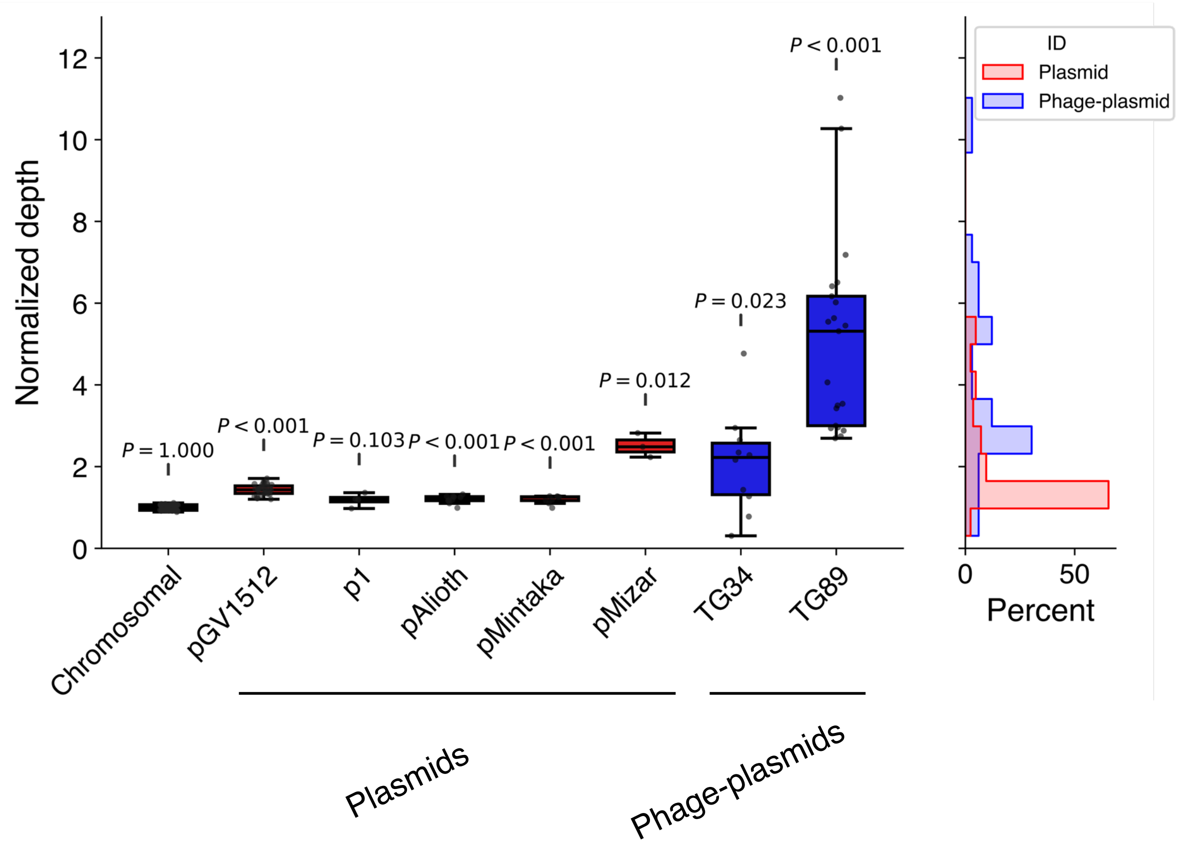


**Figure S28** Illumina short reads of the 31 *V. crassostreae* strains containing phage-plasmids of TG34 and TG89 were remapped against their scaffolded assemblies to infer their copy number in the bacterial cell. For each strain, the depth of read coverage against each variety of contig was normalized to the mean coverage against chromosomes 1 and 2. Significance shows one-sample *t*-test against an expected coverage value of 1, to represent the expected coverage of a lysogenic prophage integrated into the host chromosome. The histogram to the right shows the distribution of coverage depth for all plasmids or all phage-plasmids.

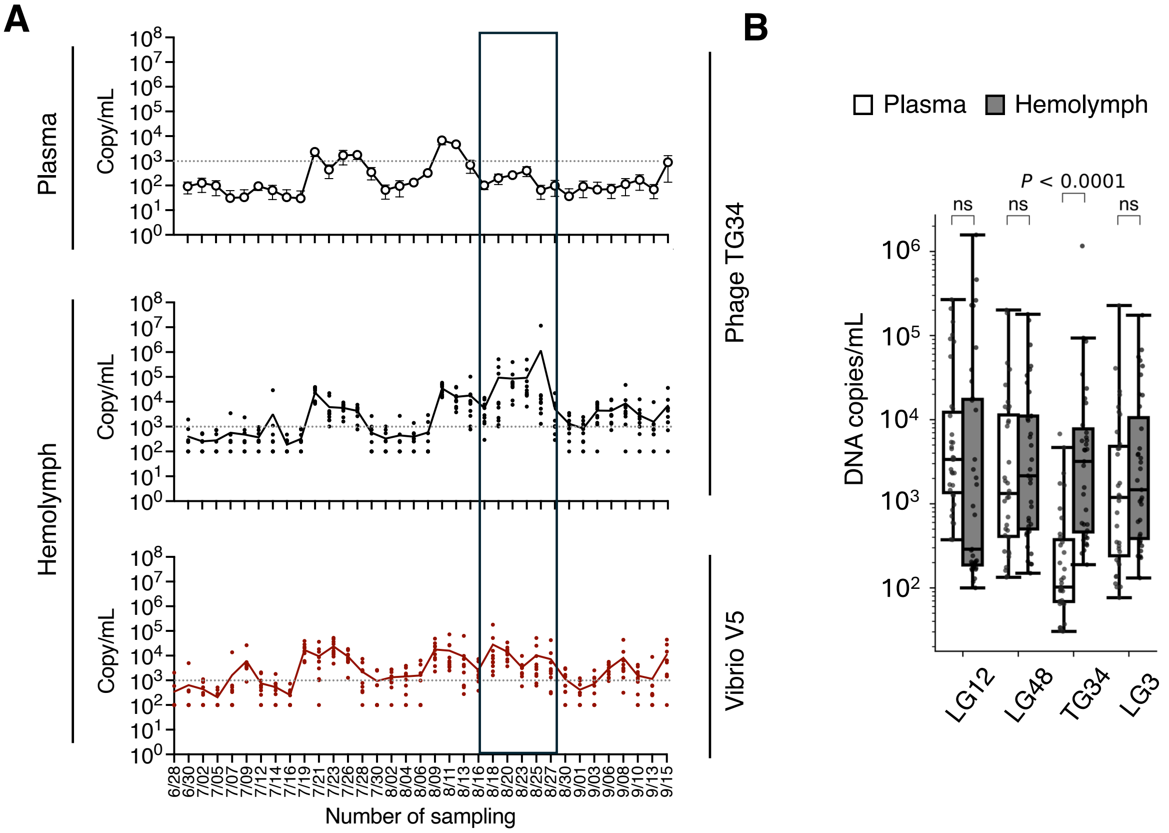


**Figure S29 Induction and lysogeny of an active temperate phage in oyster hemolymph.** **A** ddPCR quantification of the temperate phage genus TG34 and vibrio clade V5 across the sampling season, measured in plasma (pooled from 90 oysters per date, four technical replicates) and hemolymph (10 individual oysters). The dotted line marks the detection limit. Detection of phage DNA in plasma confirms activation, consistent with the recovery of lytic phages from the same genus (LG49, Figure 2A). From August 18 to 27 (inset), TG34 abundance spiked exclusively in hemolymph without a corresponding increase in V5, suggesting activation followed by lysogeny in other clades. This is consistent with the predicted host range of TG34, which includes clades V2 and V4 (Figure 5A). **B** Most virulent phages (LG3, LG12, LG48) displayed comparable abundances in plasma and hemolymph. In contrast, TG34 reached significantly higher levels in hemolymph, supporting its temperate lifestyle. Statistical significance was tested using the two-sided Brunner–Munzel test with post-hoc correction for four comparisons.

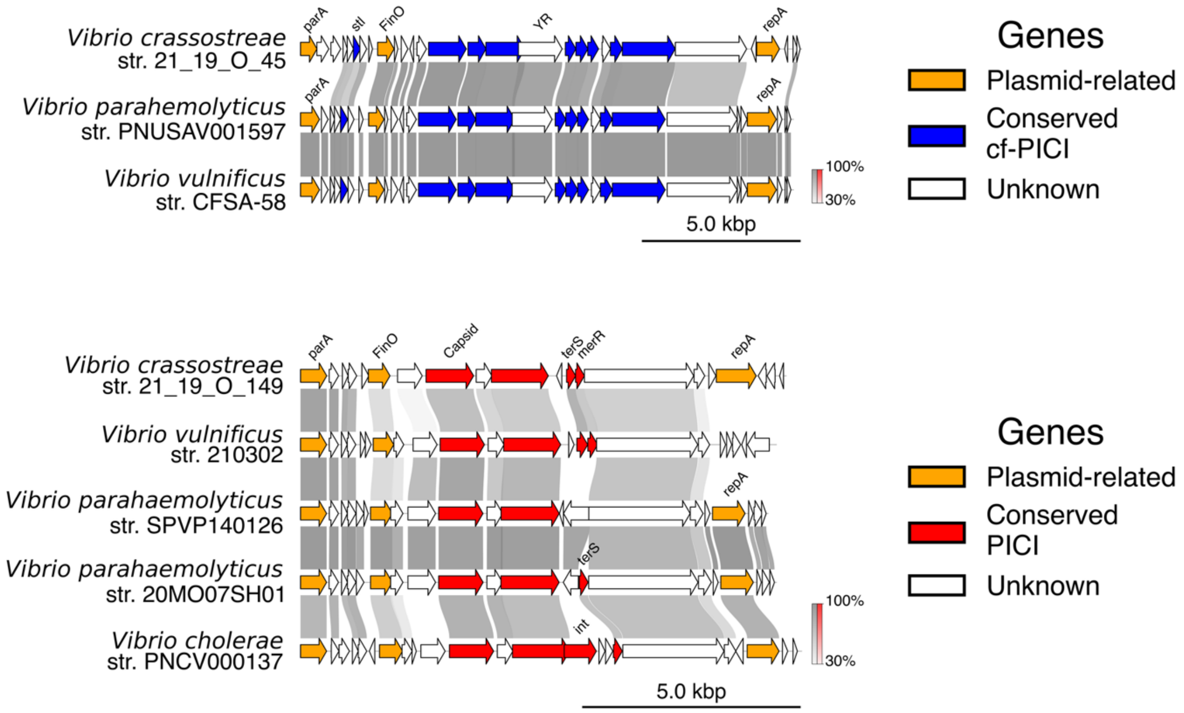


**Figure S30 Plasmid-satellites are distributed widely across *Vibrio sp.*** We screened for elements similar to the plasmid-satellites described in our dataset in public datasets by targeting assemblies with closely related capsid or parA gene sequences. Links between adjacent tracks show the percent identity of bi-directional best hits computed by MMseqs with a minimum identity of 30%. The sequence and synteny of the satellite modules are well-preserved in both the PISP and cf-PISP elements. Without exception, the homologous elements were assembled as contigs separate from the bacterial chromosomes which, together with their carriage of plasmid-like genes, strongly supports their identification as similar plasmid-satellites.

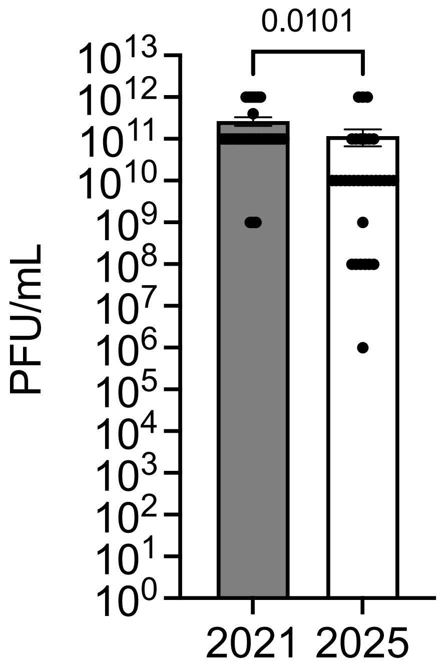


**Figure 31** Phage isolates retained viability after four years, with titers decreasing on average by ~1 log.
